# Constructing the ensemble of representative structures for a protein via neural-surrogate-guided MSA recombination

**DOI:** 10.64898/2026.01.14.699462

**Authors:** Hanyang Zhou, Hongyu Yu, Stephen S.-T. Yau, Haipeng Gong

## Abstract

Structural dynamics is essential for the functional and mechanistic illustration of proteins. Previous research attempted to generate diversified protein structures by utilizing the multiple sequence alignment (MSA), but failed to provide physically relevant representative conformations without state annotations. In this work, we propose a framework named ProCEDiS to generate a compact ensemble of representative conformations for the target protein without prior knowledge. Adopting a neural surrogate to assist the exploration of MSA recombination and integrating with AlphaFold2 to model structures, this method can automatically find high-quality, mutually dissimilar conformations for the target sequence. Parallel short-timescale molecular dynamics (MD) simulations on these structure seeds enable quick while crude free energy estimation, from which physically plausible representative states could be identified. In the benchmark on four protein systems, the ProCEDiS + MD pipeline is capable of providing valuable structural dynamics information within acceptable running time.

## Introduction

Functional and mechanistic elucidation of a protein typically requires a comprehensive characterization of its underlying conformational landscape^1–4^. Hence, an ideal computational framework should at least be capable of sampling functionally relevant regions of the overall conformational space and organizing the corresponding structural representatives in a manner consistent with thermodynamic preference. Advances of deep learning techniques have dramatically reshaped the field of protein structure prediction. Representative models like AlphaFold2^5^ and RoseTTAFold^6^ as well as their successors^7–10^ have successfully promoted the structure prediction accuracy to a level comparable to experimental structure determination by effectively exploiting the coevolutionary information encoded in the multiple sequence alignment (MSA). Unfortunately, these methods are primarily designed and optimized for single-state structure inference, while lacking the capacity to resolve the conformational polymorphism of functional proteins that are critical in maintaining and regulating the living system.

The pre-trained protein structure predictors, however, are found to be able to sample alternative conformations when perceiving perturbations to the MSA input^11^. Building on this observation, a number of algorithms have been proposed to extract the conformational diversity by decoupling or biasing the mixed evolutionary signals of multiple structural states hidden in the raw MSA^11–16^. These methods can be further classified by their MSA manipulation strategies: those relying on row-wise operations that filter, subsample or cluster homologous sequences, and those applying column-wise perturbations that alter residue-level signals through masking or mutation. The former category, exemplified by AF Cluster^15^, preserves the query sequence while separating the MSA into distinct evolutionary subsets, whereas the latter one biases local interactions by directly introducing artificial perturbations. Despite the reported enhancement in sampling alternative conformations, these methods always return a large number of unordered, often redundant structures while lacking an explicit principle to organize or rank these conformations by physical relevance, which hinders in-depth investigation and thus severely undermines their practical usefulness.

In parallel, generative modeling approaches have been developed to directly learn the conformational distributions from sequence data, shifting from signal perturbation to explicit distribution modeling^17–19^. For instance, diffusion-based frameworks like BioEmu^20^ and flow-based methods like AlphaFlow^21^ can generate a large ensemble of diverse, high-confidence structures within limited inference time. However, due to the lack of physical basis in their model design, the generated ensemble usually diverges from thermodynamic equilibrium. Moreover, automated identification of low-energy, biologically relevant structure states within such an ensemble is highly challenging, particularly in the absence of prior state annotations. Consequently, the generated structures often require additional post hoc filtering to serve as qualified starting points for more physically reliable investigations such as molecular dynamics (MD) simulations.

Indeed, a central difficulty shared by both MSA-manipulation-based approaches and generative models lies in the translation of structural diversity into a limited number of high-quality, mutually dissimilar structural representatives that are compatible with physical plausibility, in the absence of prior state annotations. Accumulating evidence suggests that polymorphic structural information is at least partially encoded within the MSA, for instance, through alternative contact patterns associated with ligand binding, membrane environment or assembly interface^15,22,23^. However, when functional states are unevenly represented across homologs, how to reliably separate and organize these signals without structural priors remains an open challenge.

To address this problem, we introduce ProCEDiS (<u>Pro</u>tein <u>C</u>onformation <u>E</u>nsemble of <u>Di</u>ssimilar <u>S</u>tructures), a discovery framework that couples evolution-guided diversity constraints with confidence- and novelty-aware selection to produce a compact set of representative conformations. This framework first applies a taxonomy- and sequence-aware MSA decomposition strategy (Taxo-seq Cluster) to generate mutually exclusive but complementary MSA sub-clusters. Then a neural surrogate is designed to guide the automatic exploration of the combinatorial space of these MSA subclusters, aiming to generate a small set of high-confidence, structurally discriminable conformational anchors under the assistance of AlphaFold2. Finally, latent-space interpolation is performed through AlphaFold2 to insert structural seeds between disconnected anchor pairs, ensuring the internal continuity of the generated structural ensemble. By this means, ProCEDiS automatically outputs a compact set of high-confidence, mutually dissimilar conformational representatives for a target protein, making the heterogeneous conformational landscape explorable with affordable computational resources. Parallel short-timescale MD simulations initiated from these seed structures enable quick while crude construction of the low-dimensional free energy landscape, from which dominant basins could be identified and ranked, with representative conformations available for mechanistic investigation and interpretation. Evaluation of this combined pipeline (ProCEDiS + MD) on four benchmark systems of versatile conformational heterogeneity patterns demonstrates that our method provides a practical route from sequence to structural dynamics, bridging MSA-based diversity extraction and physics-based sampling to systematically explore the functionally relevant protein conformational space.

## Results

### Overview of ProCEDiS

Taking AlphaFold2 (with the same parameter weights but implemented with OpenFold^24^) as the structure generator, ProCEDiS automatically outputs a compact set of high-confidence, structurally dissimilar conformation seeds for a target protein through two sequential steps: Taxo-seq Cluster and neural-surrogate-guided search (NSGS) (Figure 1a & b). After latent-space structure interpolation, the downstream workflow further relaxes and organizes the identified structure seeds by parallel short-timescale MD simulations and crude free energy estimation (Figure 1c). Overall, the ProCEDiS + MD pipeline allows automated identification of energetically stable conformational states of the target protein purely based on its sequence and MSA information.

**Figure 1.**
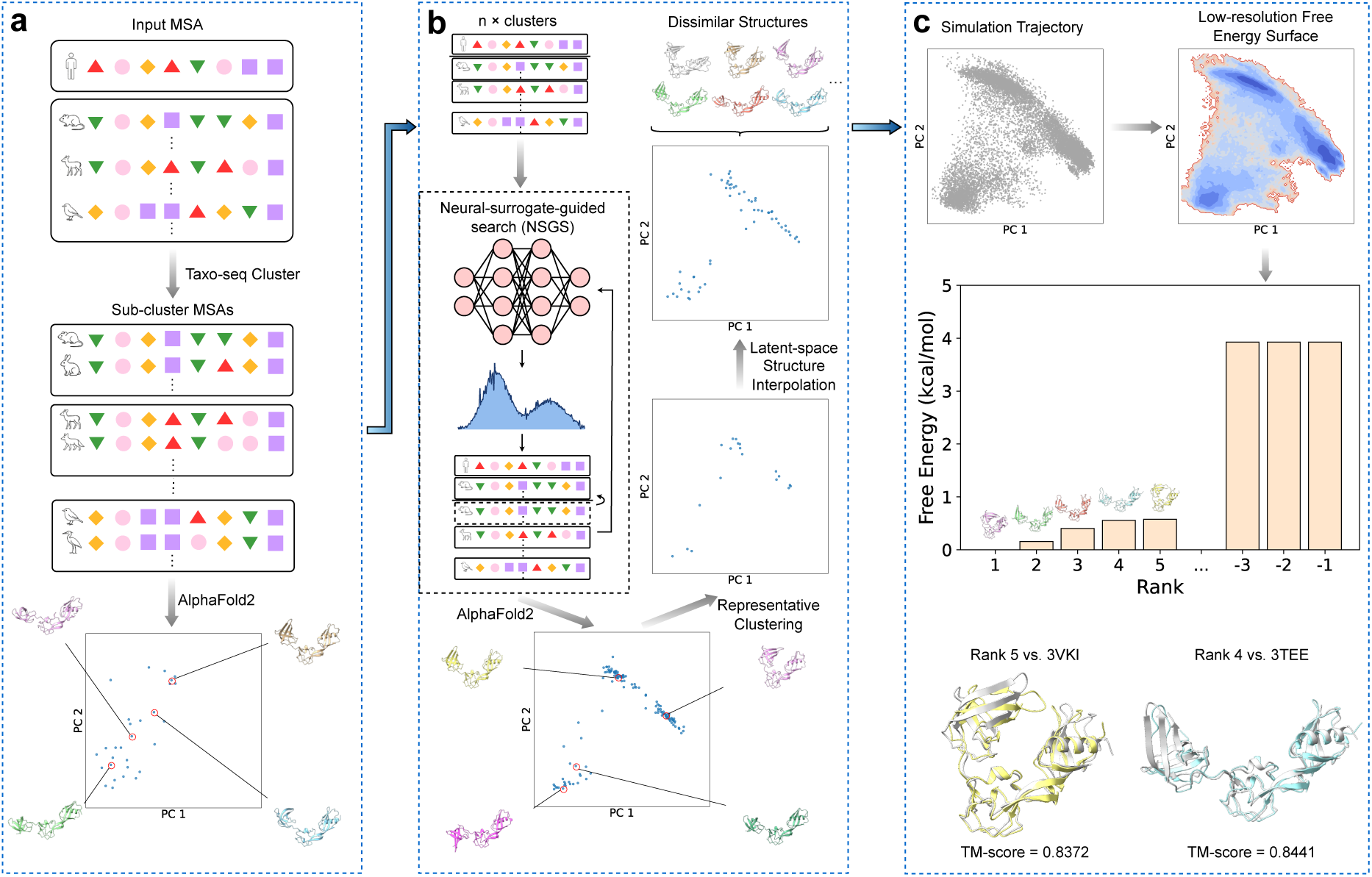
Overview of ProCEDiS. ProCEDiS constructs a compact set of conformational seeds for the target protein without prior knowledge, and can be coupled to low-cost MD simulations (ProCEDiS + MD) for crude free energy estimation. **a)** Starting from a full MSA of the query sequence, Taxo-seq Cluster generates complementary MSA sub-clusters that preserve evolutionary diversity under controlled depth. These sub-clusters are used as direct inputs for AlphaFold2 inference or as action units in the subsequent combinatorial exploration. **b)** NSGS explores the combinatorial space of MSA sub-clusters by estimating the expected structural novelty and modeling quality of candidate states using a continually updated neural surrogate. Qualified MSA subsets are passed to AlphaFold2 for structure modeling. The generated structures with satisfactory novelty and quality are added to the evolving structure pool. **c)** The ProCEDiS + MD pipeline. ProCEDiS outputs a compact set of mutually dissimilar conformational seeds by pool deduplication and latent-space structure interpolation. These seeds can be further relaxed and organized through parallel short-timescale MD simulations to construct a crude free energy estimation, which enables automated basin ranking and representative selection. In this schematic presentation, identified structural representatives (colored in green or purple) consistent with experimentally determined structures (colored in silver) are shown in structure alignment.

### Taxonomy awareness improves MSA clustering

For each target protein, we obtained mutually exclusive MSA sub-clusters through Taxo-seq Cluster, which utilizes both sequence and taxonomy relationships for information decomposition, and then generated the corresponding protein structures using AlphaFold2 (Figure 1a). In comparison to AF Cluster that relies on sequence similarity alone, Taxo-seq Cluster additionally leverages taxonomy awareness to improve the intra-cluster information purity. When evaluated on a benchmark of 77 proteins with experimentally resolved alternative conformations (*i.e.* inter-state TM-score < 0.7, see Supplementary Table 1), this amendment not only enhances the success rate of covering distinctive conformational states by AlphaFold2-folded structures (Supplementary Figure 1), but also allows comparably reliable structure modeling using MSAs of significantly reduced depths (Supplementary Figure 2). After this processing, all MSA sub-clusters compose the sequence pool, awaiting the subsequent surrogate-assisted exploration of MSA recombination (Figure 1b), while only structures of high quality (*i.e.* mean pLDDT ≥ 70) are incorporated into the structure pool for structural novelty evaluation. Overall, our clustering strategy effectively constrains the action space of MSA recombination to the simple combination of MSA sub-clusters, while retaining the complementary evolutionary signals essential for structural diversity detection.

### NSGS effectively explores the MSA recombination

To explore the combinatorial space of MSA sub-clusters while maintaining structure modeling confidence, we formulated the MSA recombination as a sequential decision process and solved it using a neural surrogate (Figure 2a). The overall process is composed of two sequential phases: a search phase and an update phase. In each episode of the search phase, the agent incrementally constructs an MSA state by selecting MSA sub-clusters from the available sequence pool, aiming to find the MSA subsets that can produce high-quality, novel protein structures. To avoid the high computational cost of repeated structure generation by AlphaFold2 in this framework, we designed a neural surrogate to quickly assess the quality (in pLDDT) of the structure corresponding to a candidate MSA state and its structural deviation (in root mean square distance (RMSD)) from those in the current structure pool purely based on the MSA information (Figure 2b). This surrogate then determines the action probabilities, intrinsically prioritizing MSA states that are likely to yield conformations having sufficient structural novelty relative to previously accepted members in the structure pool. Qualified MSA states suggested by the neural surrogate are then passed to AlphaFold2 for structure prediction, and new structure candidates with confirmed structural novelty and modeling confidence (in pLDDT) are absorbed into the structure pool. In the update phase, the same neural surrogate is conversely used to assist the identification of MSA states that are expected to improve the quality of structure candidates in the pool, and the AlphaFold2-folded structures with confirmed enhancement in modeling confidence will substitute the corresponding pool members, followed by pool deduplication. Throughout this study, structural novelty assessment and pool deduplication are implemented using a length-dependent C*_α_*RMSD threshold *τ* (Equation 4), which defines the default granularity for structure pool maintenance and seed construction but is freely tunable by users to balance compactness and redundancy of the conformational ensemble.

**Figure 2.**
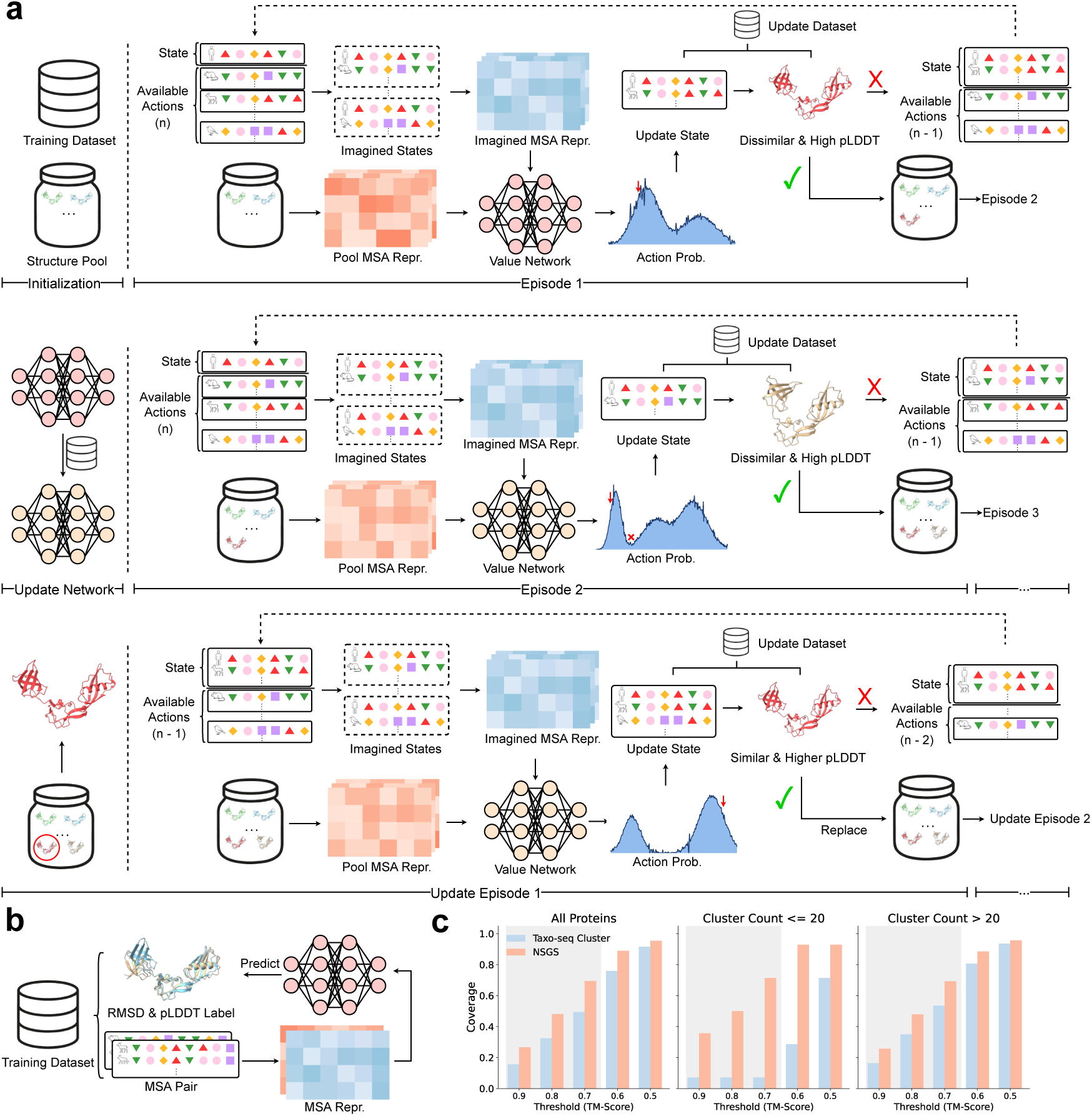
NSGS enables efficient exploration of MSA recombination. **a)** Schematic of the NSGS procedure. An initial structure pool is constructed from structures predicted based on individual MSA sub-clusters, filtered by modeling confidence and deduplicated by structural novelty. At the beginning of each episode, the MSA state contains the query sequence only. In the search phase (upper panel), a neural surrogate evaluates candidate MSA sub-clusters in combination with the current state and assigns action probabilities at each step. Candidate MSAs are inferred by AlphaFold2 for structure generation and then evaluated for structural novelty as well as modeling confidence. An MSA state is deemed successful if the predicted structure is of high-confidence and structurally novel relative to the current structure pool. Successful states expand the pool, while unsuccessful states proceed to the next step until termination. With the proceeding of the search phase (middle panel), the action probabilities change abruptly with the expansion of the structure pool, while the neural surrogate is continually updated due to its persistent learning aim. In the update phase (lower panel), the same neural surrogate is utilized to find MSA states that can produce structures of high resemblance but with improved modeling confidence to replace existing structures in the pool. **b)** Architecture of the neural surrogate. One-hot encodings of MSA pairs are embedded using a multilayer perceptron (MLP), and their difference is decoded to predict the RMSD between the corresponding structures generated by AlphaFold2 as well as the modeling confidence. **c)** Coverage analysis on the benchmark of 77 proteins with experimentally validated alternative conformations. The structure ensembles updated after NSGS (red) show improved coverage over the distinctive states than those produced by Taxo-seq Cluster alone (blue), with the largest gains in systems with no more than 20 available MSA sub-clusters.

During the exploration of MSA recombination, the probability distribution of actions is expected to change abruptly with the expansion of the structure pool in the search phase, due to the discrete alteration of targets for structure novelty assessment (Figure 2a), which obviously hinders the gradient-descent-based policy optimization in sophisticated reinforcement learning (RL) frameworks such as PPO^25^ (Supplementary Figure 8). In contrast, benefited from the simple surrogate-determined policy adopted in our NSGS approach, the neural surrogate maintains a persistent learning aim (*e.g.*, predicting structural dissimilarity from a pair of MSAs). This network is pre-trained using the plenty of MSA sub-clusters and corresponding AlphaFold2-folded structures derived from Taxo-seq Cluster, and is continually updated during the intensive exploration of MSA recombination. These characteristics jointly ensure the smooth model learning with pool expansion in the search phase (Supplementary Figure 8). Moreover, the same network is simply applied in the update phase to improve the modeling quality of pool members, without changing its learning aim. Ablation studies have confirmed the positive contribution of our surrogate network in improving the coverage of experimentally determined conformational states as well as the model quality of the derived structure ensemble (Supplementary Figure 3).

We evaluated the contribution of NSGS on the same benchmark of 77 proteins with experimentally validated alternative conformations. Here, the binary coverage of each protein conformational state (Equation 1) is evaluated by a series of TM-score thresholds and then averaged over all protein states in the benchmark (Equations 2 & 3). After 100 episodes of exploration per protein, the resulting structure pools show markedly higher coverage of major conformational states than the initial pool generated from Taxo-seq Cluster alone (Figure 2c). The most pronounced improvement is observed in challenging targets with no more than 20 available MSA sub-clusters, where purely clustering-based approaches saturate quickly. Even in this constrained regime, NSGS continually identifies novel, high-confidence conformations, indicating that diversified structure information can still be extracted from an MSA of limited evolutionary variation in the absence of state labels.

### ProCEDiS achieves superior state coverage with restrained ensemble size

In the mechanistic investigation on protein dynamics, the number of candidate structures should be restrained within an upper bound, in order to facilitate the in-depth human inspection and the follow-up computational or experimental examination. To assess the applicability of our method (Taxo-seq Cluster + NSGS) against the state-of-the-art AlphaFlow and BioEmu in such practical scenarios, we quantified the coverage of protein conformational states as a function of TM-score threshold, but restricted the number of generated representative structures by an upper bound *K*, which was set to different values (*K* = 10, 20, 30 and 40). Again, the coverage metric is evaluated on all experimentally determined conformations in the benchmark of 77 proteins, where the mean TM-score between two states of the same protein reaches 0.6055, a cutoff value representing the limit of structural distinguishability in this test set. To effectively constrain the ensemble size, we computed the pairwise C*_α_* RMSD matrices for structures generated by each method, performed agglomerative hierarchical clustering based on these matrices, cut the dendrogram to obtain exactly *K* clusters, and selected a high-quality representative structure from each cluster (see Supplementary Methods S2.5 for details). This evaluation approach standardizes the number of structures presented for examination, enabling direct comparison across methods that produce different ensemble sizes. Here, TM-score is chosen for the cover-age evaluation because it provides a length-normalized similarity measure for cross-method comparison, whereas the RMSD-based threshold *τ* is used internally to control novelty and redundancy during the construction of structure pool and seeds in NSGS.

As shown in Figure 3a, ProCEDiS comprehensively achieves a higher coverage within the regime of valid distinguishability (*i.e.* TM-score ≥ 0.6055) than AlphaFlow and BioEmu as well as Taxo-seq Cluster (an improved substitute of AF Cluster) across tested ensemble sizes (see Supplementary Table 2 for the protein-wise raw data). The advantage is more pronounced at higher similarity thresholds (*e.g.*, TM-score *>* 0.8) and smaller ensemble sizes (*e.g.*, *K* = 10), indicating that ProCEDiS preferentially recovers major conformational basins rather than distributing sampling across marginal variants. These results highlight the distinction between the conventional exhaustive structure generation and the purposed detection of representative conformations in this study: unlike generative models that randomly produce a large number of structures, ProCEDiS is focused on generating a compact set of high-quality, mutually dissimilar conformational seeds, which allow in-depth human inspection as well as MD-based exploration on the functionally relevant conformational subspace with affordable computational resources.

**Figure 3.**
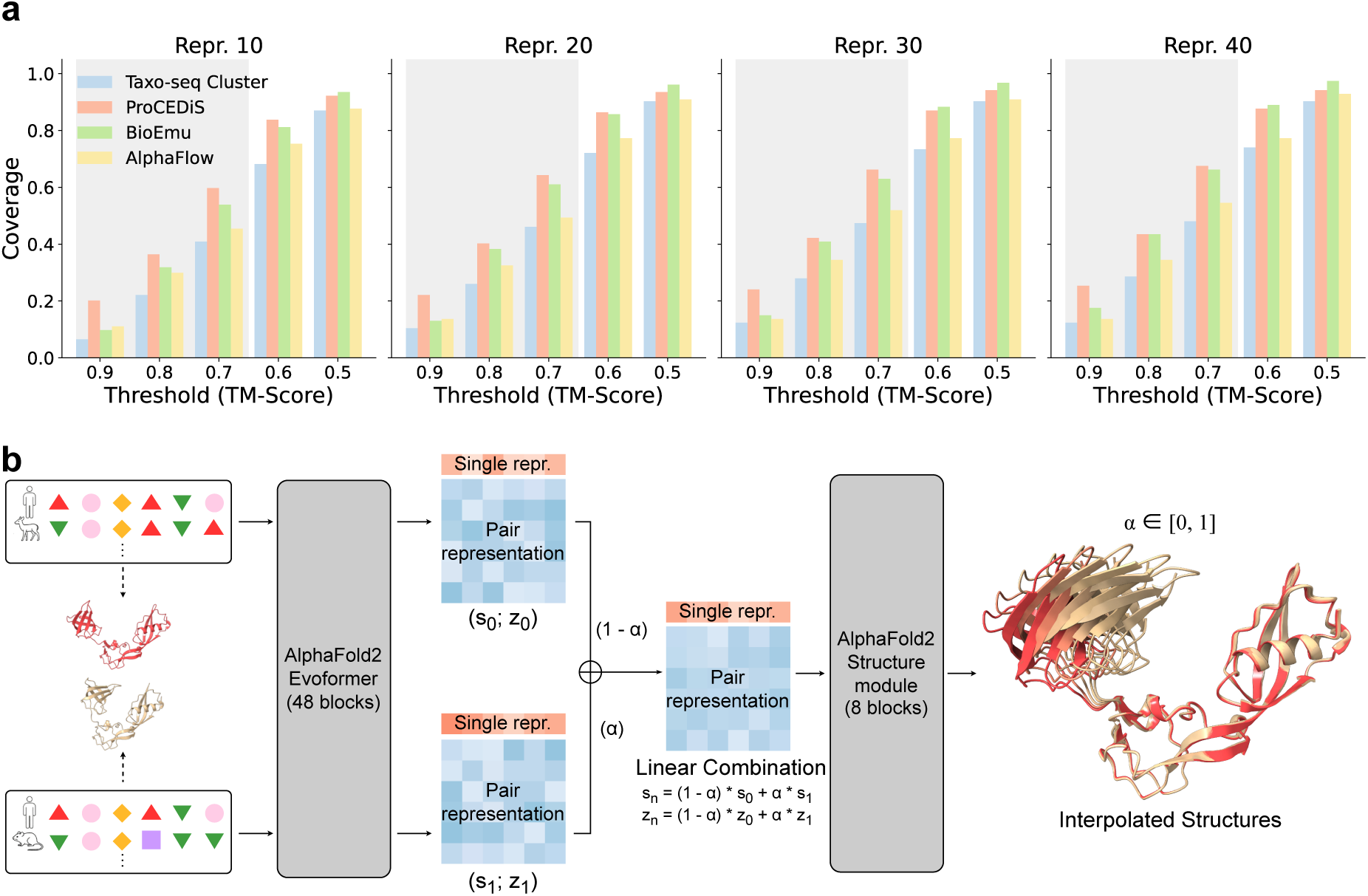
Coverage evaluation and latent-space structure interpolation. **a)** Coverage of experimentally determined conformations as a function of TM-score threshold at a series of fixed values for the number of representative structures (*K* = 10, 20, 30 and 40), evaluated on the benchmark of 77 proteins. The horizontal axis refers to the threshold (in TM-score) for coverage evaluation, whereas the vertical axis denotes the calculated coverage value. The shaded region indicates the range of valid structural distinguishability, with TM-score 0.6055 (*i.e.* the average pairwise similarity between ground-truth states). **b)** Latent-space structure interpolation. Corresponding MSA subsets of the two disconnected structures are fed into the Evoformer of AlphaFold2 to generate latent embeddings. The corresponding single and pair representations are linearly combined through a tunable factor *α* and then propagated through the Structure Module of AlphaFold2 to generate intermediate conformations that bridge the two representative states in the conformational space.

### Latent-space structure interpolation fills conformational gaps

Following the NSGS exploration, the structure pool occasionally populates isolated regions of the conformational space. To improve continuity without introducing uncontrolled diversity, we performed linear interpolation between disconnected conformation pairs in the latent space of AlphaFold2 Evoformer outputs (Figure 3b). Interpolated representations were propagated through the AlphaFold2 Structure Module to generate intermediate structures bridging distant representative states. To maintain a compact set of conformational seeds, interpolated structures were deduplicated using the same length-dependent RMSD criterion (*τ*) for seed construction in previous sections before the downstream analyses or MD simulations. Notably, interpolation was applied only after candidate anchors had been identified and validated by AlphaFold2 prediction, in order to ensure that intermediate conformations remain anchored to plausible evolutionary and structural contexts.

### Evaluation of the full ProCEDiS + MD pipeline

In this study, we evaluated the full ProCEDiS + MD pipeline on four benchmark systems spanning diverse modes of conformational heterogeneity: the periplasmic chaperone FlgA, in which conformational transitions are coupled to functional assembly^26^ (PDB IDs: 3VKI/3TEE); the conformation-driven enzyme adenylate kinase (ADK) and its two mutational derivatives^27–29^ (PDB IDs: 1AKE/1DVR/2AK3/4AKE); and two proteins explicitly excluded from the AlphaFold2 training set (PDB IDs: 7SY9/8DP2 and 8DP7/8DP6). Together, these cases stress-test whether a compact set of conformational seeds can be discovered from sequence and MSA information alone, and whether downstream physical organization can recover the interpretable conformational landscape without reliance on prior state annotations or potential training-set memorization.

#### Low-resolution FES estimation is sufficient for basin ranking in FlgA

To test whether short-timescale MD trajectories can reliably organize conformational ensembles, we first applied ProCEDiS to FlgA to generate a compact set of 57 conformational seeds, and then initiated parallel 100-ns MD simulations in explicit solvent from these seeds. After projection into the subspace of the first two principal components (PCs) through the principal component analysis (PCA), a low-resolution free energy surface (FES) could be quickly constructed from these short MD trajectories based on crude estimation of sampling frequency in each individual local bin (Figure 4a, left). This FES contains multiple stable basins, and more importantly, successfully reproduces the dominant transition between experimentally determined states (see the top path covering 3VKI and 3TEE). In contrast, the FES estimated from BioEmu-generated structures only presents one major well around the common conformational state corresponding to 3VKI, unable to describe the conformational transition process (Figure 4a, right). We failed to apply the MD-based free energy estimation on BioEmu-generated samples, because the number of candidate structure seeds created following the same structure clustering protocol exceeds 700 (Supplementary Table 3) and many of these structures have unreasonable C*_α_*-C*_α_* distances (Supplementary Table 4), both characteristics hindering efficient MD simulations with affordable computational resources. Fortunately, the MD-based free energy estimation could be successfully applied to AlphaFlow-generated structures (Supplementary Figure 4). Although the correspondingly constructed FES successfully covers the two experimentally determined structures, it contains evidently less information than the FES generated by our pipeline when projected into the same subspace.

**Figure 4.**
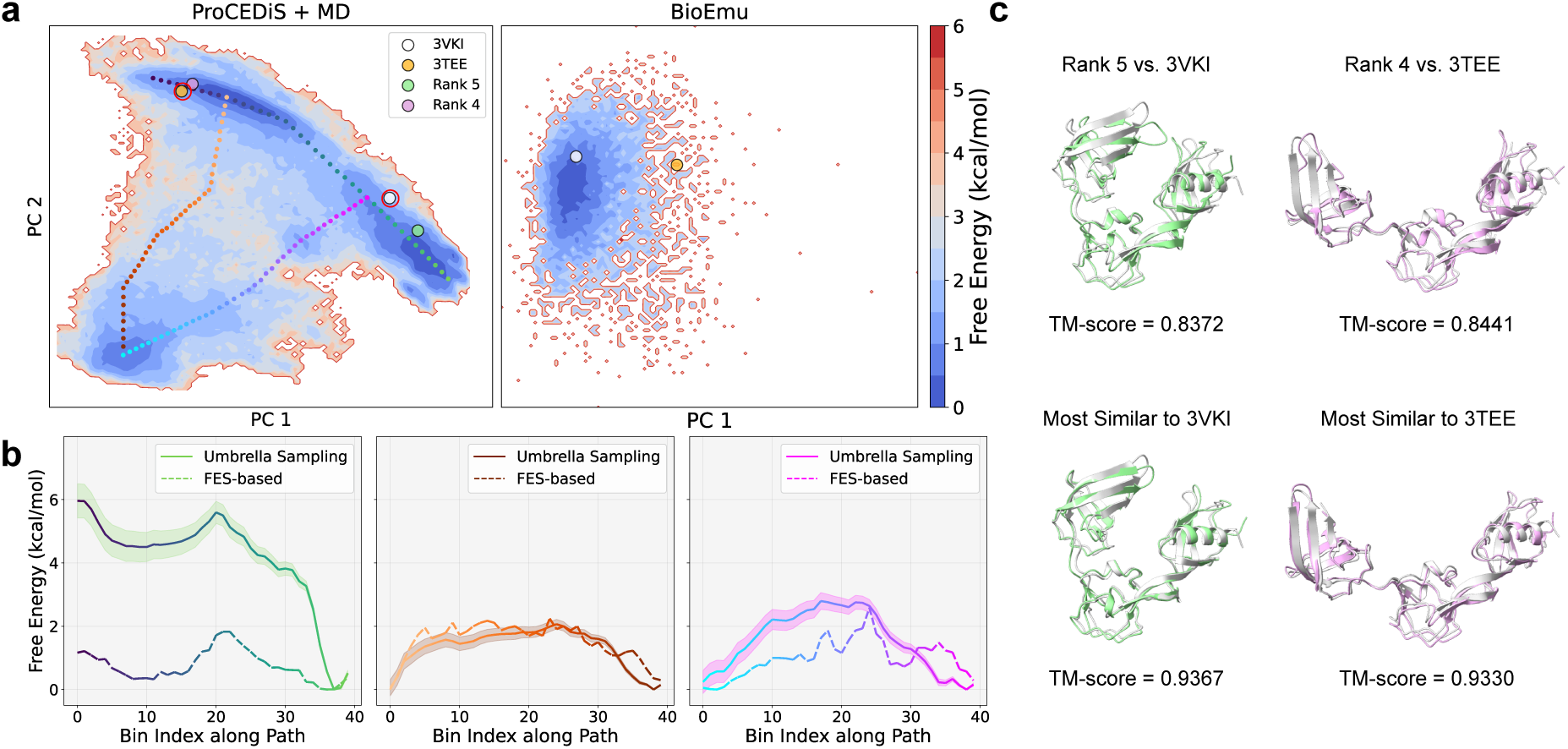
Free energy organization and basin ranking of FlgA. **a)** Low-resolution FES projected into the subspace of the first two PCs. Crude FES is constructed from conformations sampled by parallel 100-ns MD trajectories initiated from ProCEDiS seeds (ProCEDiS + MD, left) and from 10,000 structures generated by BioEmu (right), respectively. The PCA subspace is fitted independently for each ensemble. Experimentally determined reference conformations are indicated by their PDB IDs (3VKI and 3TEE) and are highlighted by red circles in the FES of ProCEDiS + MD. **b)** 1D free energy profiles along three representative transition pathways (colored identically to panel (**a**), left). Solid lines show the PMF profiles reconstructed rigorously using umbrella sampling and MBAR, with shaded regions indicating uncertainty. Dashed lines show the free energy profiles obtained directly from binning the low-resolution FES without bias reweighting. The curves are aligned by their minimum values. **c)** Representative conformations selected by automated basin identification and free-energy-based ranking. The best representatives (upper row) and the closest structures in MD trajectories (lower row) are superposed onto the ground-truth conformational states, with TM-score indicated below. Computationally generated conformations are colored in green or purple and experimental structures in silver.

To further verify the free energy landscape constructed by our pipeline, we chose three transition paths and rigorously evaluated the potentials of mean force (PMF) along them using umbrella sampling and multistate Bennett acceptance ratio (MBAR). As shown in Figure 4b, the crude 1D free energy profiles estimated by our short MD trajectories qualitatively agree with the corresponding PMF curves, with coincidence in the position of energy basins, despite the underestimation in barrier heights. Although extended simulations are expected to further improve the numerical consistence, we stopped here considering that the main purpose of this study focused on the identification of physically plausible representative structures. Through automated basin identification and free energy ranking, representative conformations with the 4th and 5th rankings successfully capture the structural natures of the two experimentally determined states (Figure 4c, upper row), although structures with higher level of resemblance exist in the MD trajectories (Figure 4c, lower row). Overall, the ProCEDiS + MD pipeline enables automated identification of physically plausible conformational states based on crude free energy estimation.

#### ProCEDiS + MD provides reasonable FES for ADK and its mutants

We applied ProCEDiS to ADK and its two mutational derivatives (P177A and Y171W) to generate the conformational seeds, followed by parallel short-timescale MD simulations and free energy analysis. Based on the convention of prior investigations on this protein^30^, MD trajectories are projected into the 2D subspace defined by two angular collective variables (*i.e.* NMP and LID domain angles), yielding structured free energy landscapes with multiple basins positioned adjacent to experimentally determined conformations (Figure 5a, left). In all of the three plots, the lowest free energy basin covers the open state conformation (corresponding to 4AKE), consistent with the knowledge that this state is preferred in the absence of substrate binding^29,31^. Furthermore, the FES presents marked changes among the three tested proteins, demonstrating the sensitivity of our computational pipeline to single point mutations in the target sequence. Intriguingly, the closed state conformation (corresponding to 1AKE) occupies a local minimum in the wild-type and P177A systems, but is destabilized by the Y171W mutation. Particularly, in comparison to the wide-type protein, the P177A and Y171W derivatives are inclined to adopt large NMP angles, consistent with the retarded diffusion observed in these systems in the absence of substrate binding^32^. After the automated basin identification and free energy sorting, in all of the three systems, both open and closed states could be captured by the limited number (set to 20 by default) of representative structures, whereas the intermediate state corresponding to 1DVR is successfully captured in the wild-type and P177A systems (Figure 5b). Albeit absent in the representative structures, the remaining state (corresponding to 2AK3) is still sampled by MD trajectories in all of the three systems (Supplementary Figure 5).

**Figure 5.**
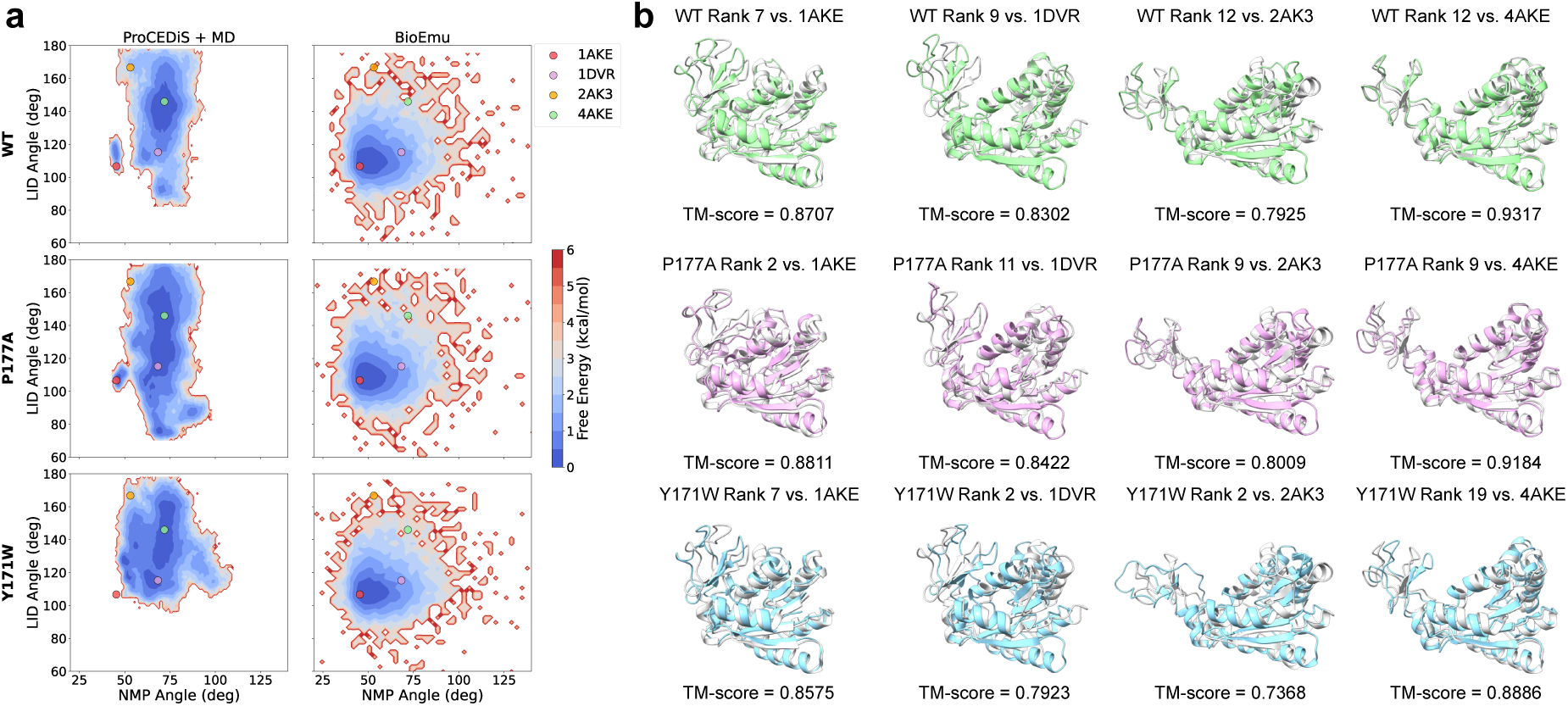
Free energy organization of ADK and its mutants. **a)** Low-resolution FES is plotted in the 2D subspace defined by the NMP and LID domain angles for the wild-type ADK (top), the P177A derivative (middle), and the Y171W derivative (bottom), respectively. Results of the ProCEDiS + MD pipeline and BioEmu are shown in the left and right panels, respectively. The four experimentally determined structures of ADK are also projected into the space, denoted by colored dots. **b)** Representative conformations selected by automated basin identification and free energy ranking. For each sequence, four representative structures corresponding to classical ADK states are superposed on their respective experimental structures, with TM-score indicated below.

In contrast, the projection of BioEmu-generated structures presents an invariant unimodal distribution for all three sequences, with the single basin overlapped with the unstable closed state conformation (Figure 5a, right). This observation implies that the lack of physical foundations in the nowadays state-of-the-art protein generative models is unlikely to be solved by simply fine-tuning over a limited, unrepresented groups of MD trajectories. Similar to the preceding section, MD-based free energy estimation is not applicable to the BioEmu samples, but is feasible to the structures generated by AlphaFlow. However, when projected into the same 2D subspace defined by NMP and LID angles, the FES of AlphaFlow + MD covers a remarkably smaller region than that of ProCEDiS + MD (Supplementary Figure 6), unable to describe the complete structural transition network for this protein.

#### FES calculation generalizes to proteins outside the AlphaFold2 training set

To assess the generalizability of our method beyond the AlphaFold2 training distribution, we applied ProCEDiS to two proteins explicitly excluded from the AlphaFold2 training set to generate conformational seeds, followed by downstream short-timescale MD simulations and free energy analysis. In the 7SY9 system, the global PCA projection directly forms a continuous FES containing multiple low free energy basins (Figure 6a), and the two experimentally determined conformational states are well described by the representative conformations derived through automated basin recognition and ranking (Figure 6b). In the 8DP7 system, although the global PCA projection presents two isolated conformational clusters, local PCA projection within the major cluster resolves a finer-scale free-energy structure with distinctive basins that are obscured in the global projection (Supplementary Figure 7). The final FES derived through such a hierarchical analysis also exhibits abundant thermodynamic information (Figure 6c), enabling the successful coverage of experimentally determined conformational states by a limited number of representative conformations automatically derived from basin identification and selection (Figure 6d). Overall, these results demonstrate that our method does not rely on training-set memorization, but instead discovers and organizes the conformational space through effectively integrating the evolutionary information and the downstream physics-based investigation.

**Figure 6.**
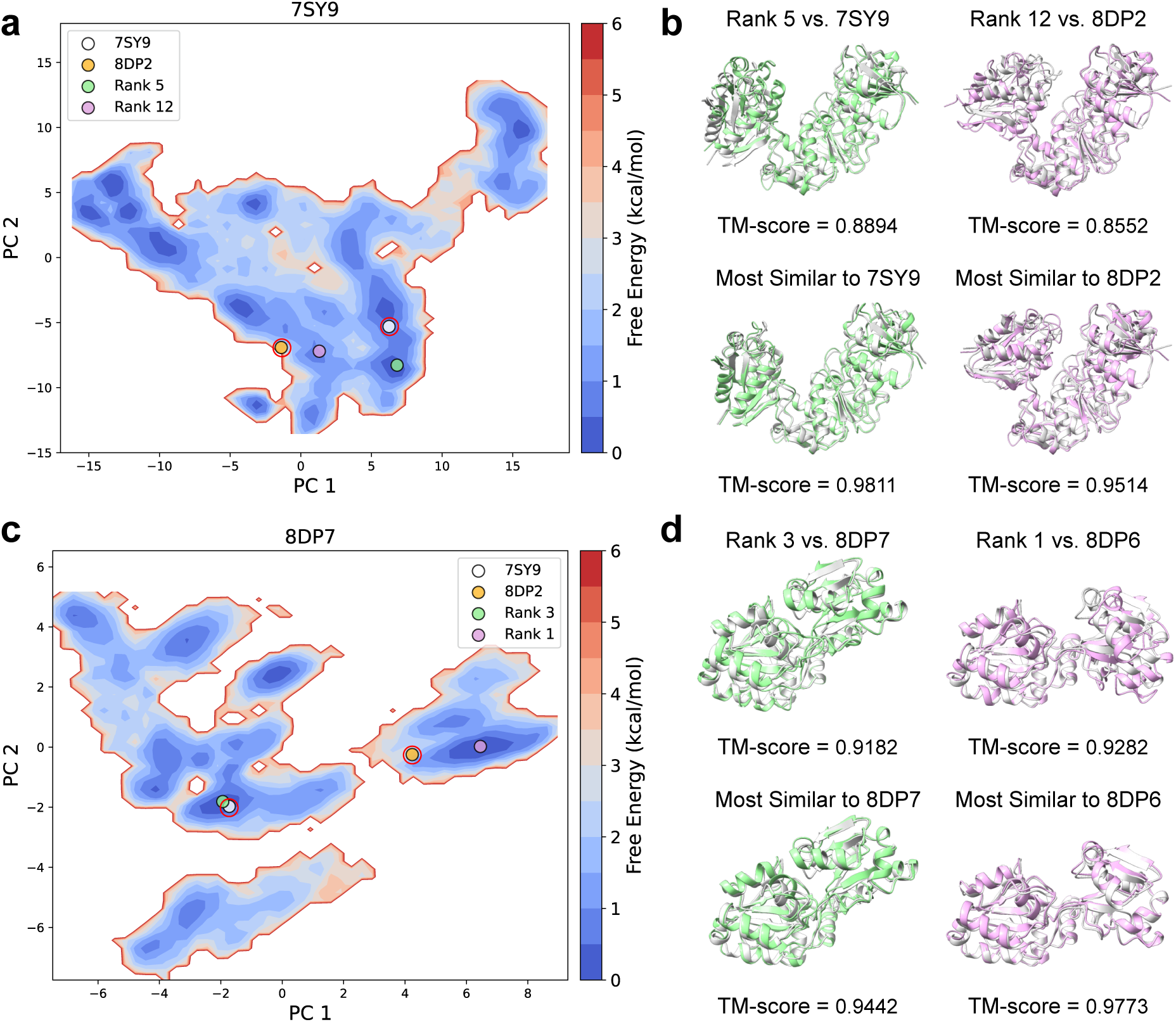
Free energy analysis for proteins outside the AlphaFold2 training set. **a)** FES of the 7SY9 system. Experimentally determined reference conformations are indicated by their PDB IDs (7SY9 and 8DP2) and are highlighted by red circles. **b)** Structural superposition of representative conformations selected by free-energy-based basin ranking (upper row) and candidate structures with high resemblance in MD trajectories (lower row) against the corresponding experimental structures for the 7SY9 system. Computationally derived conformations are shown in green or pink and reference structures in silver, with TM-score indicated below. **c)** FES of the 8DP7 system after the projection into local PC basis. Experimentally determined reference conformations are indicated by their PDB IDs (8DP7 and 8DP6) and highlighted by red circles. **d)** Structural superposition of representative conformations for the 8DP7 system, following the convention of panel (**b**).

### Running time analysis

We evaluated the running time of our pipeline on the FlgA target using 4 NVIDIA RTX 4090 GPUs (Supplementary Table 5). For this protein, ProCEDiS consumes about 4 GPU hours, most of which is taken in the NSGS exploration. This consumption is moderately higher than generative models like BioEmu and AlphaFlow (1-2 GPU hours). Taking the parallel short-timescale MD simulations in consideration, the overall ProCEDiS + MD pipeline costs slightly longer than two days. Considering that our pipeline can not only automatically identify physically relevant conformational states in the absence of any prior annotation but also provide abundant thermodynamic information, this cost is fully acceptable.

## Discussion

In this work, we introduce ProCEDiS, a framework that integrates evolutionary signal recombination, surrogate-assisted exploration and short-timescale MD simulations for the systematic exploration and organization of protein conformational space. Unlike existing approaches that emphasize large-scale structure generation, ProCEDiS is designed to identify a compact, representational ensemble that preferentially recovers major conformational basins without requiring prior state knowledge. By prioritizing fixed-budget screening over exhaustive generation, ProCEDiS targets the compact, operationally useful ensemble for mechanistic interrogation and downstream simulations.

A central contribution of this study is the explicit separation between conformational *discovery* and *organization*, which clarifies what must be learned from evolutionary data vs. what can be imposed by downstream physics-based investigation. This separation yields an operational benefit: ProCEDiS outputs a compact, high-confidence seed set that is immediately explorable, while ProCEDiS+MD provides a physics-grounded layer to relax, consolidate and rank these candidates. In practice, parallel short-timescale trajectories supply a consistent organizing signal, enabling automated basin identification in the low-dimensional FES and eliminating reliance on post hoc filtering.

Across diverse benchmark systems, our pipeline consistently demonstrates that limited and targeted MD sampling is sufficient to recover functionally relevant conformational heterogeneity. In FlgA, the low-resolution FES derived from parallel short-timescale trajectories reproduces dominant transition trends observed in the prohibitive umbrella sampling and MBAR calculations, enabling reliable basin ranking despite the limited sampling timescale. In ADK and its mutants, structured free energy landscapes emerge naturally within the biologically meaningful collective-variable space, allowing automated identification of multiple classical conformational states and mutation-induced redistribution of basin populations without explicit structural priors. Importantly, similar performance is observed for proteins explicitly excluded from the AlphaFold2 training set, indicating that our method achieves the goal not by simply memorizing the training-set information.

A key conceptual distinction highlighted by our analyses is that satisfactory structures derived by conformational generation are not necessarily suitable as seeds for MD simulations. Generative models such as AlphaFlow and BioEmu are optimized to sample broad structural distributions and excel at producing large ensembles with high apparent diversity. However, after redundancy reduction and confidence-based filtering, AlphaFlow-derived ensembles often collapse to a small number of usable structures, limiting effective conformational coverage. Conversely, BioEmu-generated ensembles typically retain substantial backbone geometric variability even after filtering. Such variablity makes it more difficult to compress an ensemble into an explorable seed set (Supplementary Table 3) and often requires additional correction on the local geometry corruption before MD seeding (Supplementary Table 4). These limitations do not reflect deficiencies in generative modeling per se, but rather expose a methodological mismatch between structure generation and seed selection for physics-based simulations. Effective MD seeding requires a balance among structural diversity, physical plausibility, and ensemble compactness — criteria that are not explicitly optimized in current generative frameworks. By contrast, our pipeline is intentionally designed to consider seed suitability: surrogate-assisted search prioritizes discriminable conformations under fixed budgets, short-timescale MD simulations enforce physical relaxation, and basin-based ranking selects representative structures that are both stable and interpretable. Moreover, this design enables ProCEDiS-derived ensembles to serve directly as non-redundant, reliable starting points for downstream simulations without extensive post hoc filtering/correction.

Several limitations of the current framework should be acknowledged. The accuracy of the low-resolution FES is constrained by the duration and number of short-timescale MD trajectories, and systems with slow or rare conformational transitions may require additional seeding or extended sampling. In addition, the surrogate-assisted search may inherit biases from the underlying structure prediction model. Future extensions could incorporate enhanced sampling techniques, experimental restraints, and/or alternative predictors to further refine basin ranking. Nevertheless, because the primary goal of ProCEDiS is to provide operationally useful MD seeds rather than quantitatively precise free energy estimates, these limitations do not diminish its practical applicability.

In summary, ProCEDiS reframes multi-conformation exploration as a problem of automated conformational screening for physics-based simulation, rather than exhaustive structure generation. By combining evolutionary information, machine learning-guided search and physics-based relaxation, the framework provides a scalable and interpretable route to identifying functionally relevant conformational ensembles that are directly compatible with MD workflows, establishing a robust foundation for downstream biophysical analysis, protein engineering and drug discovery.

## Methods

### Benchmark dataset of proteins with alternative conformations

To evaluate the ability of different methods to recover conformational heterogeneity, we curated a benchmark of proteins with experimentally resolved alternative conformations from the dataset compiled by Bryant *et al.*^33^. Proteins longer than 600 residues were excluded to ensure computational tractability. To focus on systems exhibiting substantial conformational differences, only protein pairs with inter-state TM-scores < 0.7 were retained. In addition, protein pairs with length differences > 20% (relative to the shorter sequence) were excluded. After filtering, the final benchmark consisted of 77 protein systems with comparable sequence lengths and pronounced conformational divergence (Supplementary Table 1).

### Coverage of conformational states under a TM-score hit criterion

For each benchmark target, we quantify whether a predicted ensemble recovers the experimentally determined alternative conformations using a TM-score–based hit criterion. Throughout this work, all TM-scores were computed with Biotite^34,35^. Let the set of reference conformations be R = {*R*_1_*, …, R_M_* } (in our benchmark, *M* = 2), and let S denote the set of predicted structures produced by a method.

For a TM-score threshold *t* ∈ (0, 1], we define an indicator of whether reference conformation *R_m_* is covered by the ensemble:

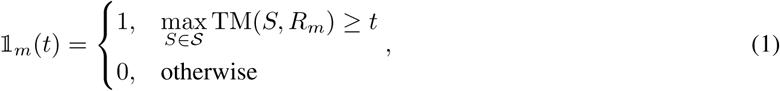

where TM(·, ·) denotes the TM-score after optimal structural alignment.

The per-target coverage at threshold *t* is then defined as the fraction of reference conformations that are hit:

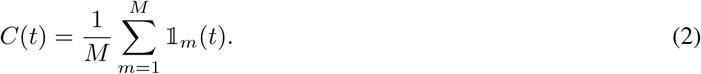

To report dataset-level performance, we average coverage across all targets:

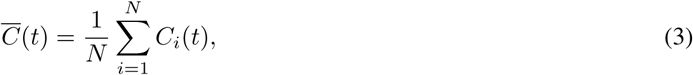

where *N* is the number of benchmark proteins and *C_i_*(*t*) is the coverage for target *i*.

### Representative selection for the ensemble-size constraining

To enable the evaluation on fixed-size ensembles (Figure 3a), we constructed representative ensembles of exactly *K* structures per target and method (*K* = 10, 20, 30 or 40). For each method, we first applied the same confidence filtering used elsewhere (mean pLDDT ≥ 70 when available), computed an all-vs-all C*_α_*-RMSD matrix over the remaining structures, and performed complete-linkage agglomerative hierarchical clustering on this distance matrix. We then cut the dendrogram to obtain exactly *K* clusters and selected one representative per cluster (highest pLDDT when available; otherwise the medoid). Full details are provided in Supplementary Methods S2.5.

### MSA generation

MSAs were generated using HHblits from the HH-suite^36^. Each query sequence was searched against the UniRef30 database^37^ for three iterations using an E-value cutoff of 0.001. During profile construction, sequences were filtered based on sequence similarity and alignment coverage to reduce redundancy while maintaining alignment quality; only hits covering at least 60% of the query sequence were retained. Final MSAs were output in the A3M format and used for subsequent analyses.

### Taxo-seq Cluster and MSA downsampling

Taxonomic identifiers (TaxIDs) were parsed from MSA annotations and mapped to NCBI taxonomy lineages using ncbitax2lin. For any pair of sequences, we defined a discrete taxonomy distance based on the deepest shared lineage level using the ordered ranks: {*below species*, *species*, *genus*, *family*, *order*, *class*, *phylum*, *kingdom*, *above kingdom*}, where smaller distances indicate closer evolutionary relatedness.

We first partitioned the taxonomy-distance matrix using complete-linkage agglomerative clustering with a fixed distance threshold corresponding to a chosen lineage level (default: family-level proximity; Supplementary Methods S1). Clusters with fewer than 5 sequences as well as sequences lacking taxonomic annotations were treated as orphans.

Within each non-orphan taxonomy cluster, we further separated mixed evolutionary signals by DBSCAN clustering on simple query-aligned one-hot sequence features (flattened vectors), following the general spirit of diversity-preserving sub-MSA construction in prior MSA-clustering approaches^15^. Orphan sequences were clustered only in this feature space. All resulting sub-MSAs, including orphan sub-MSAs, were used as candidate units for downstream exploration under fixed MSA-depth budgets. Hyperparameters were fixed across targets and are reported in the accompanying configuration files and Supplementary Methods S1.

### NSGS for conformation exploration

ProCEDiS uses the NSGS procedure to explore combinations of MSA sub-clusters and to identify structurally dissimilar, high-confidence conformational anchors. Structure inference was performed with OpenFold^24^ using pre-trained AlphaFold2 parameters^5^ (AF2-parameterized inference).

We initialized a structure pool by predicting one structure per MSA sub-cluster and filtering by confidence (mean pLDDT ≥ 70). The remaining structures were deduplicated using a length-dependent C*_α_*RMSD threshold

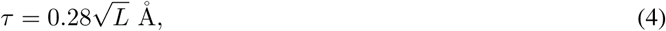

where *L* is the number of residues. This threshold defines novelty for pool expansion and controls redundancy for the seed sets reported throughout the study.

To pre-train the neural surrogate, we constructed a supervised dataset of paired MSA sub-clusters and trained the network to predict the structural RMSD between the corresponding AlphaFold2-inferred structures from differences in latent MSA representations as well as the pLDDT for an MSA state (Supplementary Methods S3). During search, each episode starts from an MSA containing only the query sequence, with MSA sub-clusters as available actions.

At each step, the neural surrogate scores candidate actions conditioned on the current state; scores are converted to sampling probabilities using a temperature-scaled Softmax function. The selected MSA sub-clusters is appended to the current state to form a candidate MSA, which is evaluated by AF2-parameterized inference. A candidate is accepted if it passes the confidence filter and is novel relative to the current pool under the RMSD threshold *τ* (the search phase). If a candidate is redundant but has higher confidence than its closest pool member, it replaces that member (the update phase). An episode terminates when an accepted structure is found or when the MSA-depth limit is reached. Hence, we treat acceptance of a novel, high-confidence structure as a terminal success signal, and use the neural surrogate as a learned heuristic to bias stochastic exploration of the combinatorial space of MSA sub-clusters. Full hyperparameters and training details are provided in Supplementary Methods S3.

### Latent-space structure interpolation for seed expansion

To fill gaps between discrete anchors identified by NSGS, we performed linear interpolation in the Evoformer latent representations produced by AF2-parameterized inference. Interpolation paths were defined only between nearest-neighbor anchors in the structural space. This restriction reduces extrapolation risk and limits the number of interpolation paths under fixed budgets. Interpolated single and pair representations were propagated through the Structure Module to obtain intermediate conformations. Paths containing any intermediate with mean pLDDT *<* 70 were discarded. The resulting structures were deduplicated under the same novelty threshold *τ* to yield a compact, high-confidence seed set. Full details are provided in Supplementary Methods S4.

### Parallel short-timescale MD simulations

Seeds produced by ProCEDiS were subjected to parallel short-timescale MD simulations for physical relaxation and downstream organization (ProCEDiS + MD). Simulations were performed with OpenMM^38^. Seeds were standardized and imaged under periodic boundary conditions using MDTraj^39^. Each system was solvated in explicit solvent with 0.15 M ionic strength, followed by energy minimization, NPT equilibration and 100-ns production simulations. Force-field choices, solvation details and simulation parameters are described in Supplementary Methods S6 & S7.

### Free energy estimation and representative structure extraction from MD trajectories

Parallel short-timescale MD trajectories initialized from ProCEDiS seeds were pooled to estimate the low-dimensional free energy landscape and to extract representative structures of physical relevance. For each trajectory, we used the OpenMM-reported potential-energy time series to automatically detect and discard an initial nonequilibrated segment, and then subsampled the remaining frames to reduce temporal correlations (MBAR timeseries utilities; Supplementary Methods S9). The resulting uncorrelated frames from all trajectories were concatenated, aligned by least squares superposition, and projected into the 2D subspace defined by the first two PCs of C*_α_*coordinates (Supplementary Methods S8) or by empirical collective variables (Supplementary Methods S10).

We discretized the projected coordinates to a regular 2D grid (typically 100 × 100 bins) to estimate the empirical density *p*(*x, y*), and computed the FES as

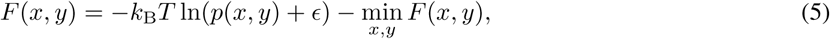

where *k*_B_ is the Boltzmann constant, *T* is the temperature, and *ɛ* is a small pseudocount for numerical stability. Representative structures were selected by iteratively ranking low-free-energy bins on the grid: we identified the current minimum-energy bin, selected the closest sampled frame to the bin center as the basin representative, masked a local neighborhood to enforce basin separation, and repeated until the desired number of basins was obtained.

### Umbrella sampling and free energy reconstruction

For selected systems, umbrella sampling simulations were performed to validate free energy trends obtained from short-timescale MD trajectories. Collective variables were defined based on PCA analysis of MD trajectories. The FES was reconstructed using MBAR^40–42^. Detailed umbrella sampling protocols and MBAR implementation are provided in Supplementary Methods S8 & S9.

## Data Availability

The dataset of proteins with experimentally determined alternative conformations and all corresponding evaluation results are available at Zenodo: https://zenodo.org/records/18231270. Atomic coordinates of MD trajectories in this work are available at the same site.

## Code Availability

All codes of the ProCEDiS pipeline, including the training scripts and model parameters, are freely accessible at GitHub repository: https://github.com/zhybio/ProCEDiS.

## Author contribution

H. G. proposed the theory and concept. H. Z. and H. G. designed the methodology. H. Z. accomplished coding, model training, performance evaluation, and data analysis. H. G. supervised the whole process. H. Y. and S. S.-T. Y. designed the protocol for taxonomy-based sequence clustering and contributed to the corresponding analysis. H. Z. and H. G. wrote the initial draft, while all authors were involved in revision. All authors agreed with the final manuscript.

## Acknowledgements

This work was supported by the Ministry of Science and Technology of China (Grant No. 2023YFF1204400 to H.G.), the National Natural Science Foundation of China (Grant No. 32171243 to H.G.), and the Beijing Frontier Research Center for Biological Structure. This work was also supported by the National Natural Science Foundation of China (Grant No. 12171275 to S. S.-T. Y.), the Tsinghua University Education Foundation (to S. S.-T. Y.), and the Beijing Natural Science Foundation (Grant No. IS25032 to S. S.-T. Y.).

## Supplementary Information

### Supplementary Methods

#### S1. Taxonomy-distance construction and the two stages of Taxo-seq Cluster

Taxonomic information was extracted from sequence annotations in the MSA using NCBI Taxonomy. Lineages were generated by converting NCBI taxdump files with ncbitax2lin (https://github.com/zyxue/ncbitax2lin). For each pair of sequences (*i, j*), we computed a discrete taxonomy distance *d_ij_* based on the lowest common taxonomic rank shared by their lineages. We used the ordered scale

*d* ∈ {0*, …,* 8} ⇐⇒ below species, species, genus, family, order, class, phylum, kingdom, above kingdom,

where smaller *d* indicates closer evolutionary relatedness (that is, a lower and more specific shared rank). The category *above kingdom* covers pairs without a meaningful shared rank under the available annotations.

##### Stage 1: taxonomy-based clustering

We first clustered the taxonomy-distance matrix using agglomerative hierarchical clustering with complete linkage (AgglomerativeClustering in scikit-learn^1^) on the pre-computed distance matrix. We did not fix the number of clusters; instead, we cut the dendrogram at a rank-level threshold. Specifically, we used a distance threshold of level + 0.5 (default level=3), which for integer-valued *d_ij_* enforces a maximum within-cluster taxonomy distance of ≤ level (that is, Family-level proximity for level=3). Clusters with fewer than 5 sequences were marked as *orphan* and processed separately in Stage 2.

##### Stage 2: sequence-feature sub-clustering within taxonomy clusters

To further separate mixed evolutionary signals within each non-orphan taxonomy cluster, we performed an additional DBSCAN sub-clustering step on simple sequence features. Each sequence was represented by a query-aligned one-hot encoding (length *L*, alphabet size 21) flattened into a vector in R^21×^*^L^*. For each taxonomy cluster, we ran DBSCAN (min_samples=5) on these one-hot feature vectors and selected the neighborhood radius eps by sweeping a fixed, target-independent set of candidate values (as specified in the released configuration). We chose the eps that maximized the number of *valid* sub-clusters (ties broken by the first maximum), where validity requires each reported sub-cluster to contain at least 5 sequences. This heuristic is used to encourage sequence diversity in the resulting sub-MSAs without using any structural labels or downstream evaluation signals.

##### Orphan handling

Sequences in orphan taxonomy clusters (size *<* 5) were pooled and clustered using the same DBSCAN procedure on one-hot sequence features, producing orphan sub-MSAs.

All hyperparameters were held fixed across targets and are reported in the accompanying configuration files / code release.

#### S2. Structure pool initialization, confidence filtering and structural deduplication

All structure predictions were performed using OpenFold^2^ with pre-trained AlphaFold2 parameters^3^. For each MSA sub-cluster, a predicted structure was generated and C*_α_*coordinates were extracted. Pairwise C*_α_*RMSDs were computed to build a structural distance matrix used for deduplication. Structures were filtered by prediction confidence (pLDDT ≥ 70) prior to pool inclusion.

Structural deduplication was performed by agglomerative hierarchical clustering with complete linkage on the C*_α_* RMSD matrix and then cutting the dendrogram at the length-dependent threshold *τ* (defined in Equation 4). The highest-pLDDT structure in each cluster was retained.

##### S2.5. Representative selection for the ensemble-size constraining (Figure 3a)

To compare the coverage of conformational state using the same number of representative structures, we constructed fixed-size representative ensembles of size *K* (with *K* = 10, 20, 30 or 40) per target for each method. For a given method, we first computed an all-vs-all C*_α_*-RMSD matrix over its generated structures after confidence filtering (pLDDT ≥ 70 when applicable). We then performed agglomerative hierarchical clustering with complete linkage on this RMSD matrix and cut the dendrogram to obtain exactly *K* clusters. One representative structure was selected per cluster, yielding exactly *K* representatives used for downstream coverage evaluation.

Unless otherwise noted, the representative for each cluster was chosen as the structure with the highest pLDDT within that cluster (for methods that output pLDDT), and as the medoid (minimum average RMSD to other members in the cluster) for methods without a directly comparable confidence score.

We emphasize that this fixed-*K* selection procedure is used only for the fixed-budget comparisons in Figure 3a. It is distinct from the length-dependent RMSD threshold *τ* used elsewhere in ProCEDiS for pool deduplication and NSGS novelty checking, where *τ* serves as a default, user-tunable granularity parameter for defining structural non-redundancy (Equation 4).

#### S3. Neural-surrogate-guided search (NSGS)

We formulated the MSA recombination as a sequential decision process. At the beginning of each episode, the state is initialized as an MSA state containing only the query sequence. Available actions are the set of MSA sub-clusters (and optionally orphan sub-MSAs). The state is represented by a binary vector indicating which MSA units have been selected.

A neural surrogate was trained to estimate (i) the structural dissimilarity between structures predicted from two candidate MSA states and (ii) the confidence of a candidate MSA state. Training samples consist of pairs of MSA embeddings, with the ground-truth label assigned as the C*_α_*RMSD between the corresponding predicted structures. In addition, we supervised the surrogate to predict the mean pLDDT associated with each MSA state, and optimized a weighted sum of MSE losses for the RMSD and pLDDT predictions.

At each step, the current state was combined with each available action to form a set of imagined next states. Embeddings of imagined states were computed and evaluated against the current structure pool P, yielding predicted RMSDs to each pool entry. For each action, we used the minimum predicted RMSD across the pool as the action score, *i.e.*, an estimate of the distance to the nearest existing conformation (larger values indicate higher expected novelty). We further applied a surrogate confidence gate by discarding actions whose predicted mean pLDDT was below *δ*. Action scores were temperature-scaled and converted to a sampling distribution using the Softmax function. For shallow MSAs (below a preset depth), actions were sampled uniformly at random to encourage exploration; otherwise, actions were sampled from the Softmax distribution.

The sampled action was appended to the current state to form a new MSA, which was passed to AF2-parameterized inference (OpenFold with pre-trained AlphaFold2 parameters) for structure prediction. A predicted conformation was deemed successful if it satisfied the confidence criterion (mean pLDDT ≥ *δ*) and the novelty criterion

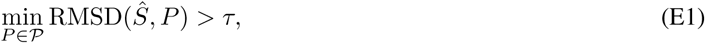

where *Ŝ* denotes the newly predicted structure, is the current structure pool, and *τ* is the threshold defined in Equation 4 (we use 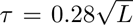). If the structure pool is empty, success reduces to the confidence criterion (mean pLDDT ≥ *δ*). If success was achieved, the predicted structure was added to the pool and the episode terminated. Otherwise, the episode continued until either success or reaching a pre-defined maximum MSA depth.

When a newly predicted structure fell within *τ* of an existing pool entry, we applied a conservative replacement rule: if there existed *exactly one* pool item with RMSD *< τ* and the new structure had higher pLDDT, we replaced that pool entry. With multiple parallel players, newly appended candidates were merged into a shared pool by accepting a candidate only if its minimum RMSD to the current shared pool exceeded *τ*.

In addition to novelty-driven search episodes, we ran update episodes that start from an anchor sampled from the current structure pool and aim to locally refine existing pool entries. Concretely, an anchor state *s*_anc_ (with structure *S*anc and mean pLDDT *p*_anc_) was sampled from P with a soft preference toward lower-confidence entries. Starting from *s*_anc_, we sampled actions to prioritize higher predicted pLDDT, while softly discouraging steps whose predicted RMSD to the anchor exceeded a local radius *d*_upd_ = 0.5*τ*. After OpenFold evaluation, the anchor entry was replaced if the new structure satisfied: (i) mean pLDDT ≥ *δ*, (ii) a minimum pLDDT improvement margin *p*_new_ *> p*_anc_ + Δ (we use Δ = 0.01), and (iii) a locality constraint RMSD(*Ŝ*new*, Ŝ*anc) ≤ *d*_upd_. Update episodes thus refine the pool locally, while preserving the novelty-driven search behavior of NSGS.

States and predicted structures encountered during both search and update were stored in an experience memory when they satisfied the confidence criterion, and were used to intermittently update the neural surrogate. Updated parameters were broadcast to all players for subsequent action selection.

##### Algorithm 1

NSGS for conformation search

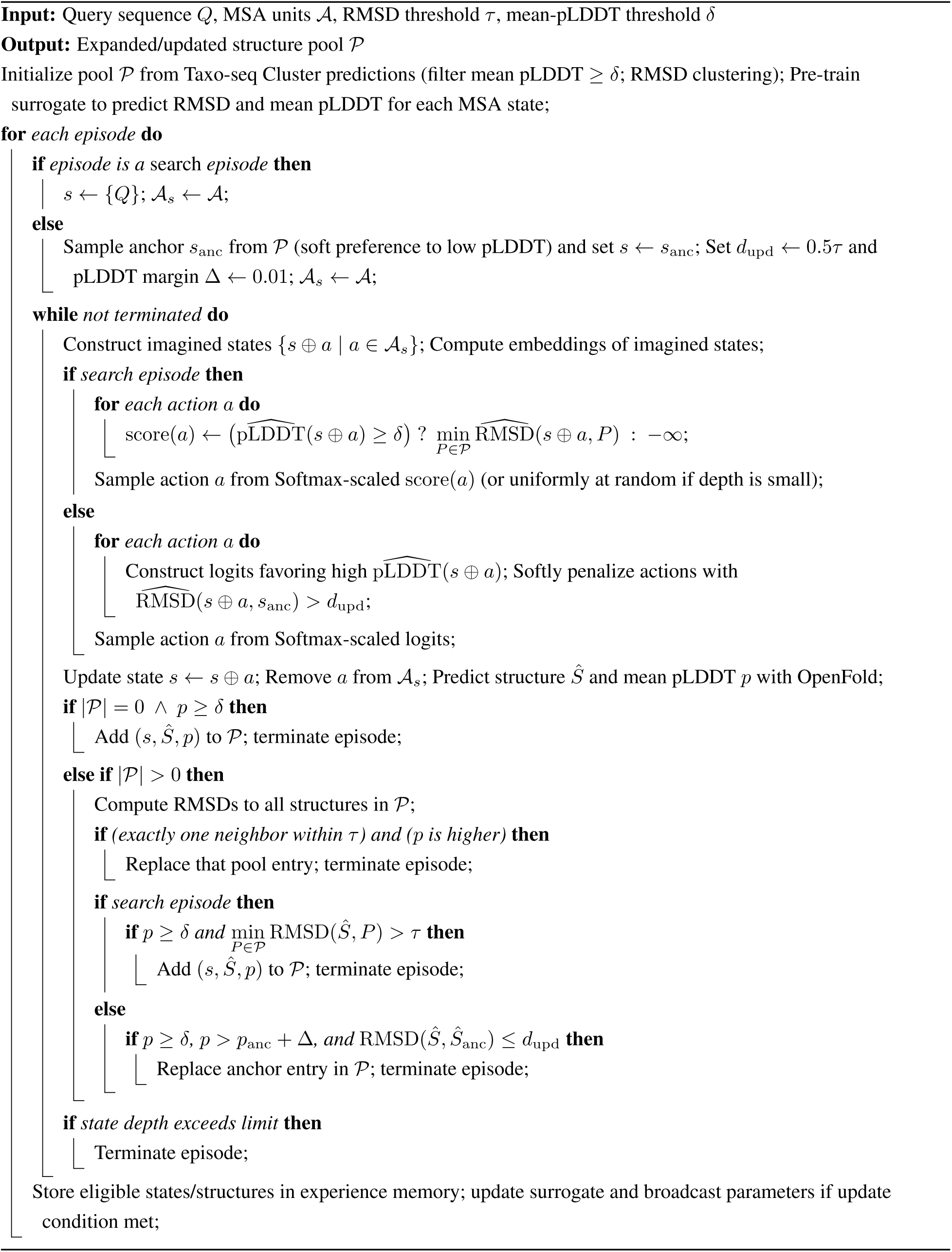

#### S4. Latent-space structure interpolation and post-processing

After the NSGS exploration, we performed an additional round of structural deduplication to account for potential structure replacement events. Only the highest-pLDDT representative in each similarity cluster was retained. Interpolation paths were defined only between each structure and its nearest neighbor in the pool, while paths with identical endpoints but opposite directions were discarded.

We linearly interpolated AF2-parameterized Evoformer outputs between endpoint conformations and fed the interpolated representations into the AF2-parameterized Structure Module (OpenFold) to generate intermediate 3D structures. Any interpolation path containing intermediate structures (excluding endpoints) with pLDDT *<* 70 was discarded. All interpolated structures were deduplicated by agglomerative clustering on the C*_α_* RMSD matrix with a distance-threshold cut at *τ* (defined in Equation 4) and then retaining the highest-pLDDT representative per cluster.

#### S5. Side-chain reconstruction for BioEmu samples

BioEmu generates structures in a backbone-frame representation, which does not contain explicit all-atom side-chain coordinates. For analyses that require all-heavy-atom models (*e.g.*, backbone-geometry and local-geometry sanity checks reported in Supplementary Table 4), we performed side-chain reconstruction following the procedure recommended in the BioEmu repository. Specifically, we used the bioemu.sidechain_relax module, which interfaces with HPacker for side-chain placement. We did not run MD equilibration; instead, we used the default setting that performs side-chain reconstruction followed by local energy minimization (*i.e.* without the optional short NVT equilibration).

Because side-chain reconstruction is computationally expensive, we applied it to a representative subset of BioEmu samples. For each target, we first clustered the 10,000 BioEmu-generated structures and selected 50 representative cluster centers (target_cluster=50). Side-chain reconstruction was then performed only on these 50 representative structures. Throughout the Supplementary Information, “BioEmu (SC)” refers to these representative structures after side-chain reconstruction, whereas “BioEmu” refers to the original backbone-frame outputs prior to side-chain reconstruction.

#### S6. MD seed preparation and system construction

All MD systems were constructed and pre-processed using OpenMM^4^ (v8.0). Each seed structure was standardized using PDBFixer (https://github.com/openmm/pdbfixer) by adding missing residues/atoms when possible and adding hydrogens at pH 7.0. All seeds were loaded in MDTraj^5^ for centering and alignment, and the maximal geometric extent across seeds was used to estimate solvation requirements under a fixed buffer distance (1.0 nm) and target ionic strength (0.15 M).

To avoid systematic difference in system size across seeds, we adopted a “fixed solvent count” strategy rather than a fixed box size. Specifically, a representative seed was solvated once under electroneutrality and 0.15 M ionic strength, and the total number of solvent molecules (water plus ions) was recorded. This total solvent count was then used as the target for all seeds. Final solvation was performed with the OpenMM Modeller module using addSolvent(…, numAdded=*N*_solv_) (OpenMM v8.0), while allowing the box vectors to adjust accordingly, where *N*_solv_ denotes the target total number of solvent molecules (water plus ions) estimated from the representative seed. We verified that the final solvent-molecule count (water+ions) matched *N*_solv_ across all seeds by topology-based counting. Systems were parameterized with the AMBER14 force field (amber14-all.xml) and a TIP3P-family water model. The finalized systems were saved in the PDBx/mmCIF format for production simulations.

#### S7. Short-timescale molecular dynamics (MD) simulations

All MD simulations were performed using OpenMM with standardized settings. Nonbonded interactions were treated with particle mesh Ewald (PME) electrostatics and a real-space cutoff of 1.0 nm. Bonds involving hydrogen atoms were constrained (HBonds). Hydrogen mass repartitioning was applied to set hydrogen masses to 1.5 amu, enabling a 4-fs integration timestep. We verified stable temperature/pressure control and constraint convergence under these settings, and did not observe abnormal energy drift (see code/logs in the release).

Dynamics were propagated using a Langevin middle integrator at 300 K with a friction coefficient of 1 ps^−1^. Each system was first energy-minimized, and the minimized structure was saved for record. Pressure coupling was achieved using a Monte Carlo barostat at 1 atm and 300 K. This was followed by 1 ns NPT equilibration and 100 ns NPT production. Energies, temperature, and box/thermodynamic properties (including volume and density) were recorded at fixed intervals, and trajectories were saved for downstream analyses. All simulations were executed in parallel with one GPU per system.

#### S8. Umbrella sampling simulations

Umbrella sampling was carried out using OpenMM coupled with PLUMED^6,7^. Collective variables were defined as the first two principal components (PC1 and PC2) obtained from PCA analysis on C*_α_* coordinates of short-timescale MD trajectories. In each umbrella window, a harmonic bias potential was applied:

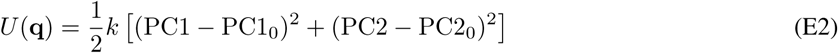

where (PC1_0_, PC2_0_) denotes the window center and *k* is the force constant. All windows were simulated independently at 300 K under NPT conditions. Simulation lengths per window, window placement, and force constants are reported in the accompanying umbrella sampling setup files / code release. During sampling, PC1, PC2, and the applied bias energy were recorded at fixed intervals.

#### S9. Free energy reconstruction using the multistate Bennett acceptance ratio (MBAR)

Free energy landscapes were reconstructed from umbrella sampling trajectories using MBAR^8–10^. For each window, time series of collective variables and bias energies were extracted from PLUMED outputs. Reduced potentials for all samples across all windows were computed by combining the unbiased potential energy with the umbrella bias term. These reduced potentials were used as MBAR inputs to estimate the unbiased probability distribution and the relative free energies. The final two-dimensional free energy surface (FES) was reported in the PC1–PC2 subspace, with the free energy reference set to the global minimum. Uncertainties were obtained from MBAR self-consistent error estimates.

#### S10. Collective variable definitions for ADK hinge-angle landscapes

For adenylate kinase (ADK), we used the conventional NMP and LID hinge-angle coordinates defined by Beckstein *et al.*^11^ as collective variables. For each trajectory frame, we computed centers of geometry (CG) of selected residue groups from C*_α_*coordinates and then evaluated inter-segment angles.

The NMP angle *θ*_NMP_ was computed from three CG points:

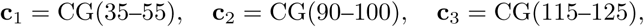

and defined as the angle between vectors **c**_1_ − **c**_2_ and **c**_3_ − **c**_2_:

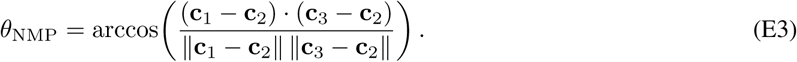

Similarly, the LID angle *θ*_LID_ was computed from

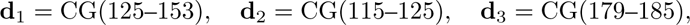

as

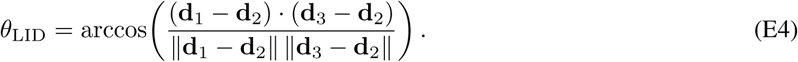

Angles were reported in degrees after conversion from radians. These two angles (*θ*_NMP_*, θ*_LID_) were then used as the 2D coordinates for grid-based FES estimation as described in **Methods**.

#### S11. Success ratio for hit-based evaluation (as implemented)

For Supplementary Figure 1, we report a “success ratio” that quantifies the fraction of method-generated structures that match the experimentally determined reference conformations above a TM-score threshold.

For each benchmark target *i*, let the two reference conformations be R*_i_* = {*R_i,a_, R_i,b_*} and let the method generate an ensemble S*_i_* = {*S_i,_*_1_*, …, S_i,ni_* } of *n_i_* predicted structures. For a TM-score threshold *t* ∈ (0, 1], we define the per-target success ratio as

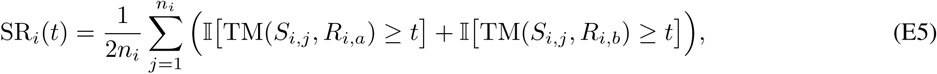

where I[·] is the indicator function (equal to 1 if the condition holds and 0 otherwise) and TM(·, ·) denotes the TM-score after optimal structural alignment. Intuitively, SR*_i_*(*t*) is the fraction of all (predicted structure, reference state) pairs whose TM-score exceeds the threshold.

We then report the benchmark-averaged success ratio across *N* targets as

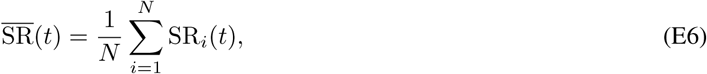

and visualize uncertainty using the standard error of the mean (s.e.m.) across targets:

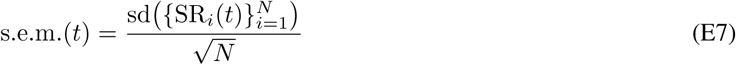

where sd(·) denotes the sample standard deviation.

### Supplementary Tables

**Supplementary Table 1.**
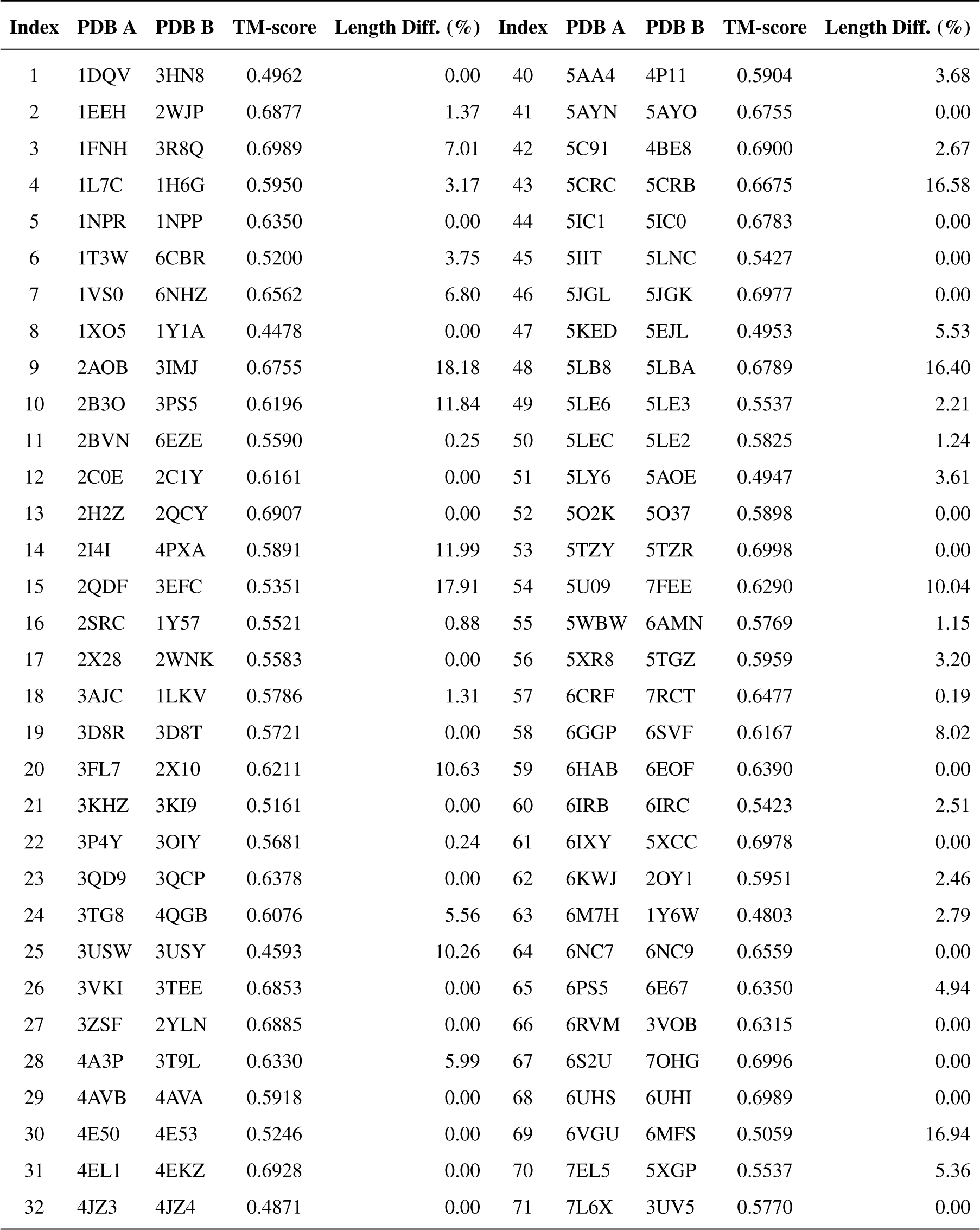
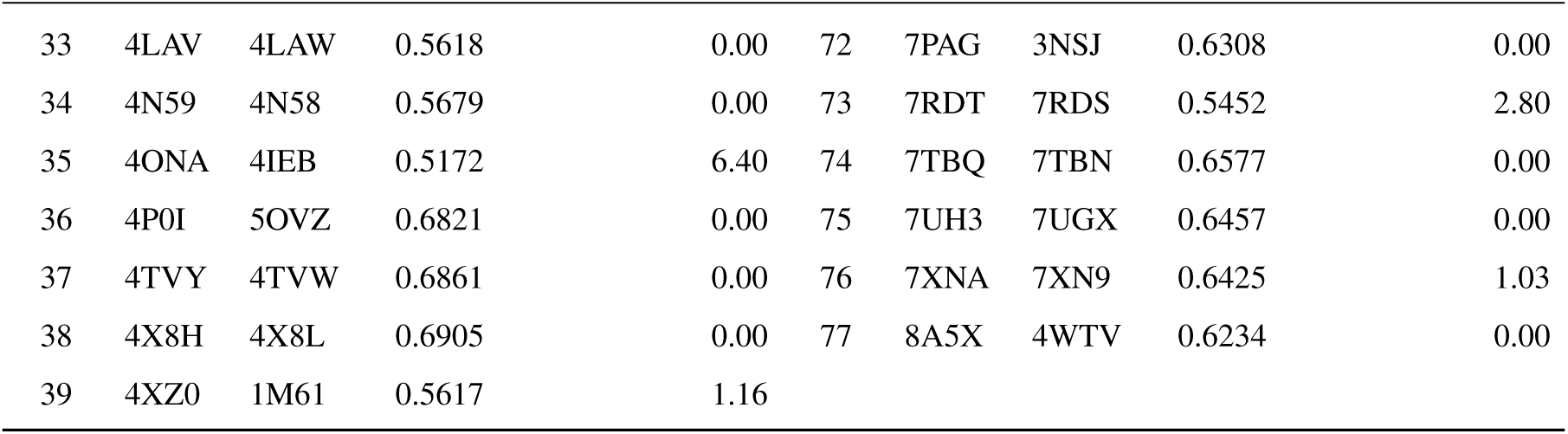
Benchmark proteins with alternative conformations. List of protein pairs (PDB A and PDB B) used to evaluate the coverage of conformational states. TM-score reports the structural similarity between the two reference conformations, and *Length Diff. (%)* denotes the relative sequence-length difference between the paired structures.

| Index | PDB A | PDB B | TM-score | Length Diff. (%) | Index | PDB A | PDB B | TM-score | Length Diff. (%) |
| --- | --- | --- | --- | --- | --- | --- | --- | --- | --- |
| 1 | 1DQV | 3HN8 | 0.4962 | 0.00 | 40 | 5AA4 | 4P11 | 0.5904 | 3.68 |
| 2 | 1EEH | 2WJP | 0.6877 | 1.37 | 41 | 5AYN | 5AYO | 0.6755 | 0.00 |
| 3 | 1FNH | 3R8Q | 0.6989 | 7.01 | 42 | 5C91 | 4BE8 | 0.6900 | 2.67 |
| 4 | 1L7C | 1H6G | 0.5950 | 3.17 | 43 | 5CRC | 5CRB | 0.6675 | 16.58 |
| 5 | 1NPR | 1NPP | 0.6350 | 0.00 | 44 | 5IC1 | 5IC0 | 0.6783 | 0.00 |
| 6 | 1T3W | 6CBR | 0.5200 | 3.75 | 45 | 5IIT | 5LNC | 0.5427 | 0.00 |
| 7 | 1VS0 | 6NHZ | 0.6562 | 6.80 | 46 | 5JGL | 5JGK | 0.6977 | 0.00 |
| 8 | 1XO5 | 1Y1A | 0.4478 | 0.00 | 47 | 5KED | 5EJL | 0.4953 | 5.53 |
| 9 | 2AOB | 3IMJ | 0.6755 | 18.18 | 48 | 5LB8 | 5LBA | 0.6789 | 16.40 |
| 10 | 2B3O | 3PS5 | 0.6196 | 11.84 | 49 | 5LE6 | 5LE3 | 0.5537 | 2.21 |
| 11 | 2BVN | 6EZE | 0.5590 | 0.25 | 50 | 5LEC | 5LE2 | 0.5825 | 1.24 |
| 12 | 2C0E | 2C1Y | 0.6161 | 0.00 | 51 | 5LY6 | 5AOE | 0.4947 | 3.61 |
| 13 | 2H2Z | 2QCY | 0.6907 | 0.00 | 52 | 5O2K | 5O37 | 0.5898 | 0.00 |
| 14 | 2I4I | 4PXA | 0.5891 | 11.99 | 53 | 5TZY | 5TZR | 0.6998 | 0.00 |
| 15 | 2QDF | 3EFC | 0.5351 | 17.91 | 54 | 5U09 | 7FEE | 0.6290 | 10.04 |
| 16 | 2SRC | 1Y57 | 0.5521 | 0.88 | 55 | 5WBW | 6AMN | 0.5769 | 1.15 |
| 17 | 2X28 | 2WNK | 0.5583 | 0.00 | 56 | 5XR8 | 5TGZ | 0.5959 | 3.20 |
| 18 | 3AJC | 1LKV | 0.5786 | 1.31 | 57 | 6CRF | 7RCT | 0.6477 | 0.19 |
| 19 | 3D8R | 3D8T | 0.5721 | 0.00 | 58 | 6GGP | 6SVF | 0.6167 | 8.02 |
| 20 | 3FL7 | 2X10 | 0.6211 | 10.63 | 59 | 6HAB | 6EOF | 0.6390 | 0.00 |
| 21 | 3KHZ | 3KI9 | 0.5161 | 0.00 | 60 | 6IRB | 6IRC | 0.5423 | 2.51 |
| 22 | 3P4Y | 3OIY | 0.5681 | 0.24 | 61 | 6IXY | 5XCC | 0.6978 | 0.00 |
| 23 | 3QD9 | 3QCP | 0.6378 | 0.00 | 62 | 6KWJ | 2OY1 | 0.5951 | 2.46 |
| 24 | 3TG8 | 4QGB | 0.6076 | 5.56 | 63 | 6M7H | 1Y6W | 0.4803 | 2.79 |
| 25 | 3USW | 3USY | 0.4593 | 10.26 | 64 | 6NC7 | 6NC9 | 0.6559 | 0.00 |
| 26 | 3VKI | 3TEE | 0.6853 | 0.00 | 65 | 6PS5 | 6E67 | 0.6350 | 4.94 |
| 27 | 3ZSF | 2YLN | 0.6885 | 0.00 | 66 | 6RVM | 3VOB | 0.6315 | 0.00 |
| 28 | 4A3P | 3T9L | 0.6330 | 5.99 | 67 | 6S2U | 7OHG | 0.6996 | 0.00 |
| 29 | 4AVB | 4AVA | 0.5918 | 0.00 | 68 | 6UHS | 6UHI | 0.6989 | 0.00 |
| 30 | 4E50 | 4E53 | 0.5246 | 0.00 | 69 | 6VGU | 6MFS | 0.5059 | 16.94 |
| 31 | 4EL1 | 4EKZ | 0.6928 | 0.00 | 70 | 7EL5 | 5XGP | 0.5537 | 5.36 |
| 32 | 4JZ3 | 4JZ4 | 0.4871 | 0.00 | 71 | 7L6X | 3UV5 | 0.5770 | 0.00 |
| 33 | 4LAV | 4LAW | 0.5618 | 0.00 | 72 | 7PAG | 3NSJ | 0.6308 | 0.00 |
| 34 | 4N59 | 4N58 | 0.5679 | 0.00 | 73 | 7RDT | 7RDS | 0.5452 | 2.80 |
| 35 | 4ONA | 4IEB | 0.5172 | 6.40 | 74 | 7TBQ | 7TBN | 0.6577 | 0.00 |
| 36 | 4P0I | 5OVZ | 0.6821 | 0.00 | 75 | 7UH3 | 7UGX | 0.6457 | 0.00 |
| 37 | 4TVY | 4TVW | 0.6861 | 0.00 | 76 | 7XNA | 7XN9 | 0.6425 | 1.03 |
| 38 | 4X8H | 4X8L | 0.6905 | 0.00 | 77 | 8A5X | 4WTV | 0.6234 | 0.00 |
| 39 | 4XZ0 | 1M61 | 0.5617 | 1.16 |  |  |  |  |  |

**Supplementary Table 2.**
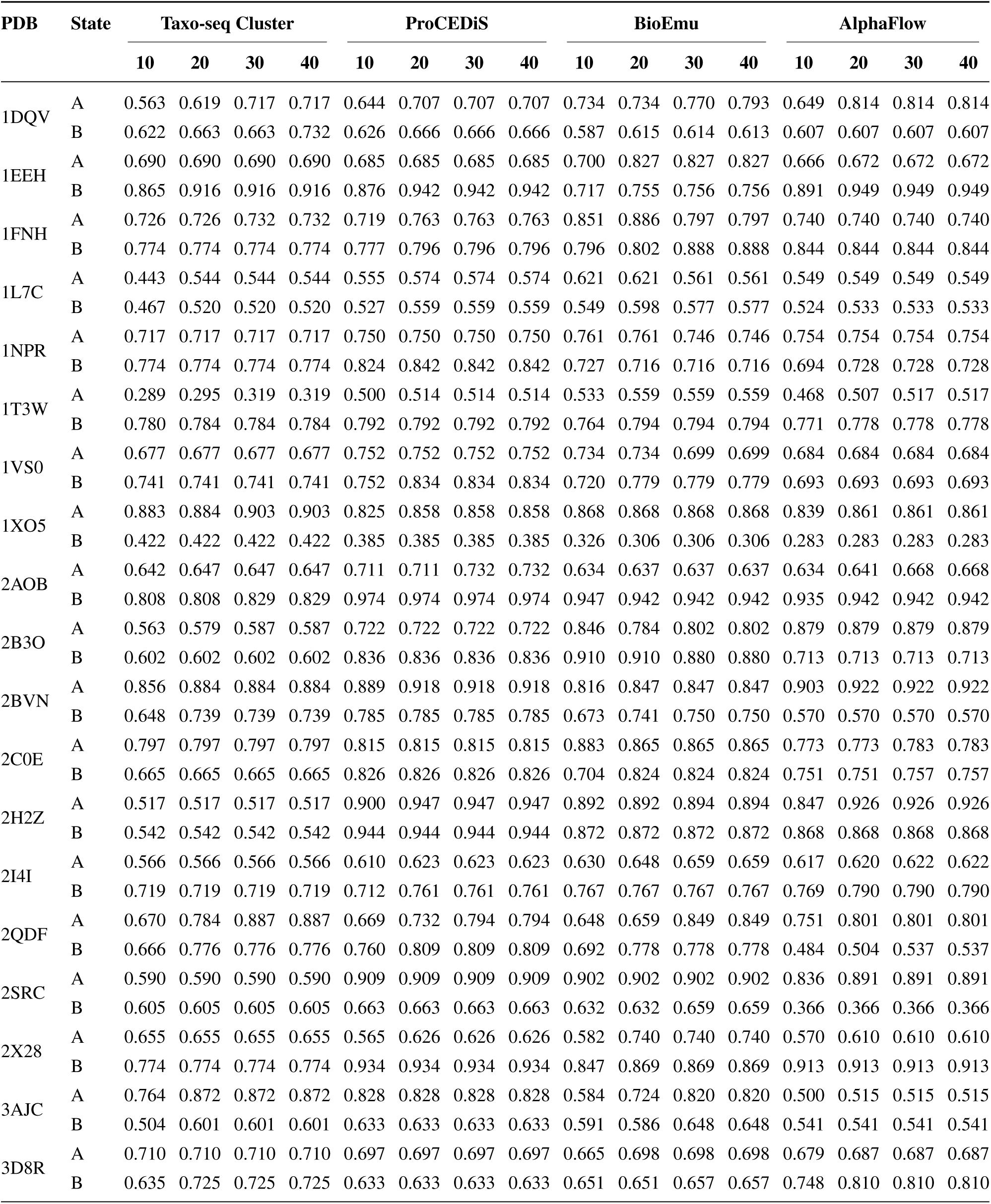
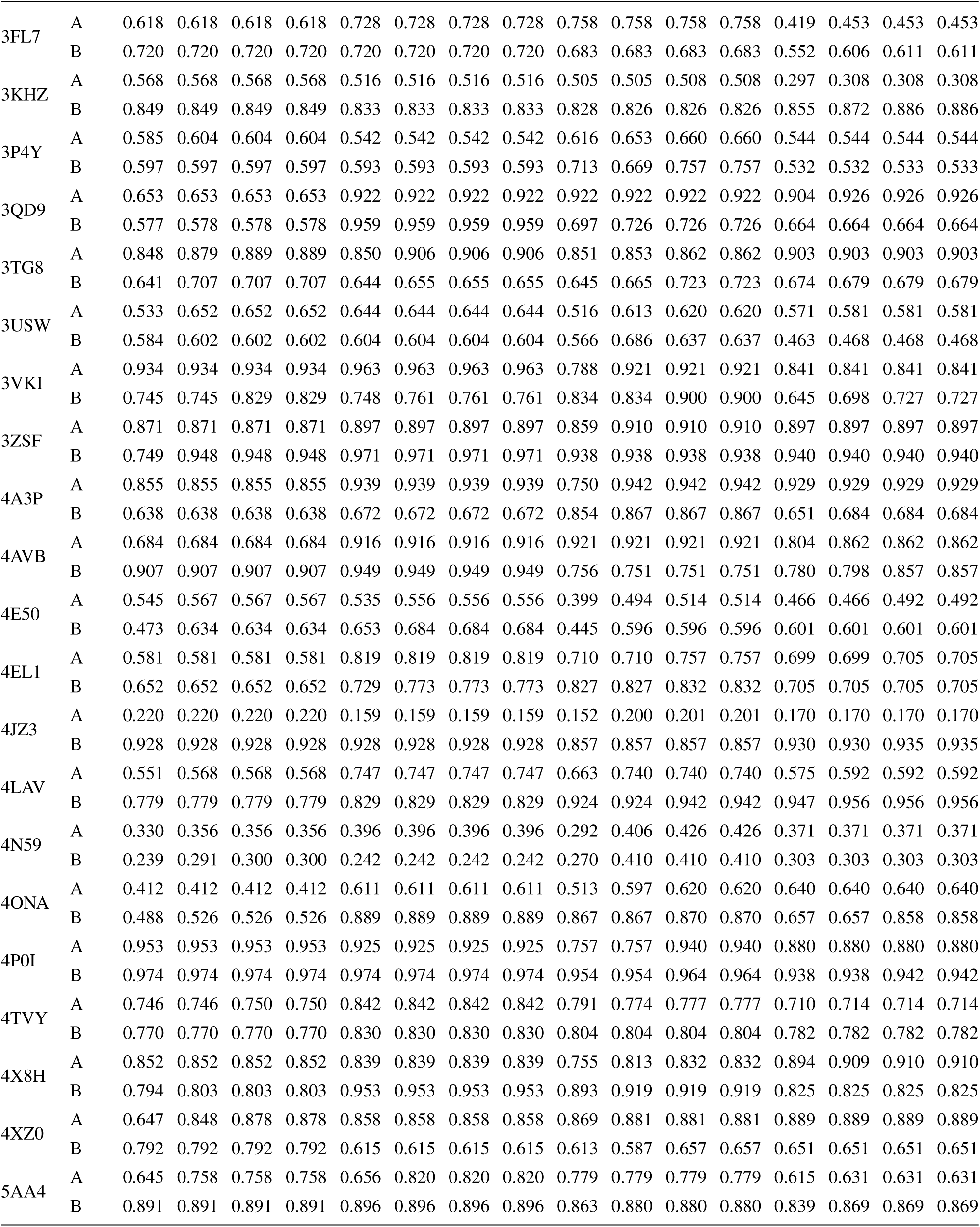
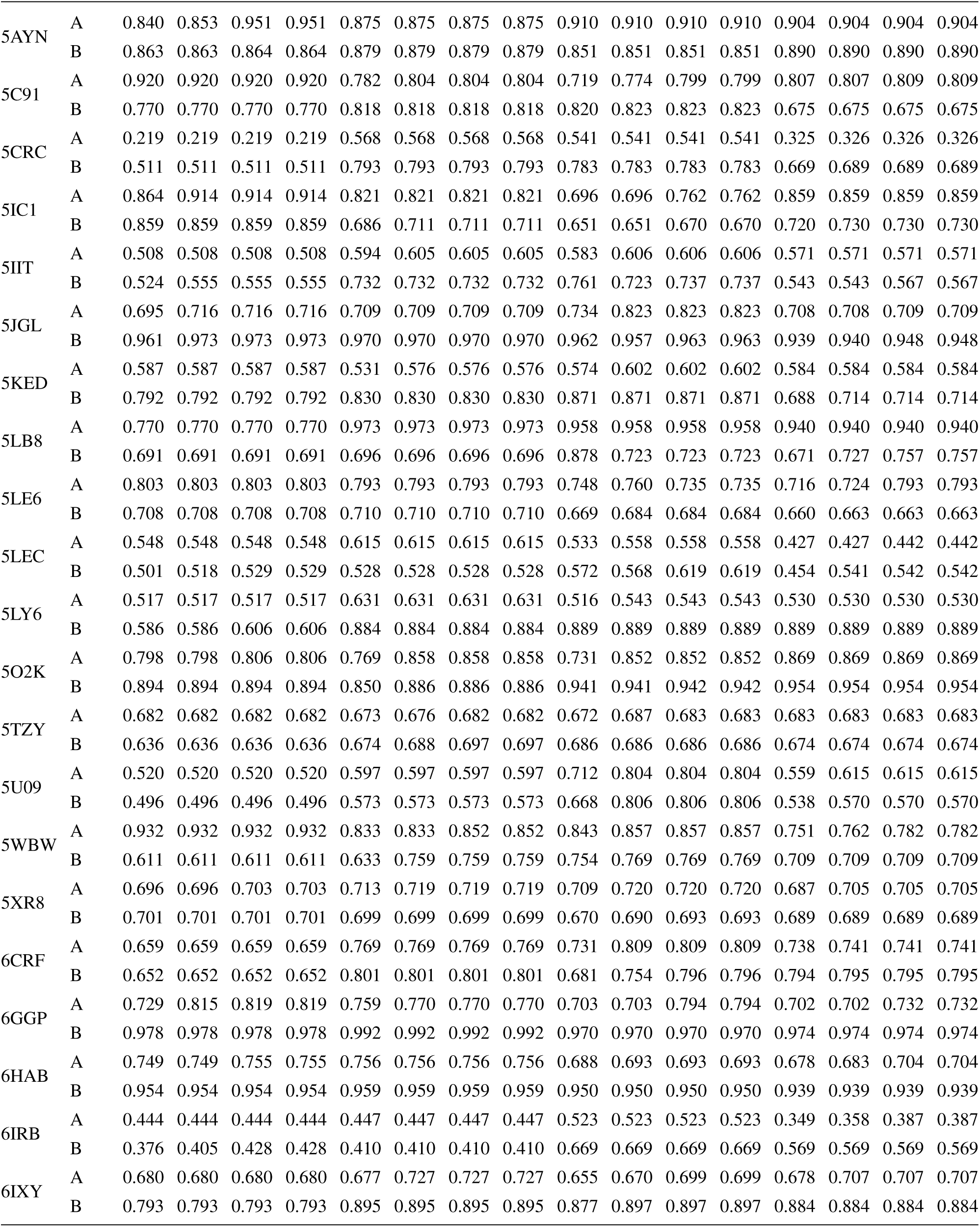
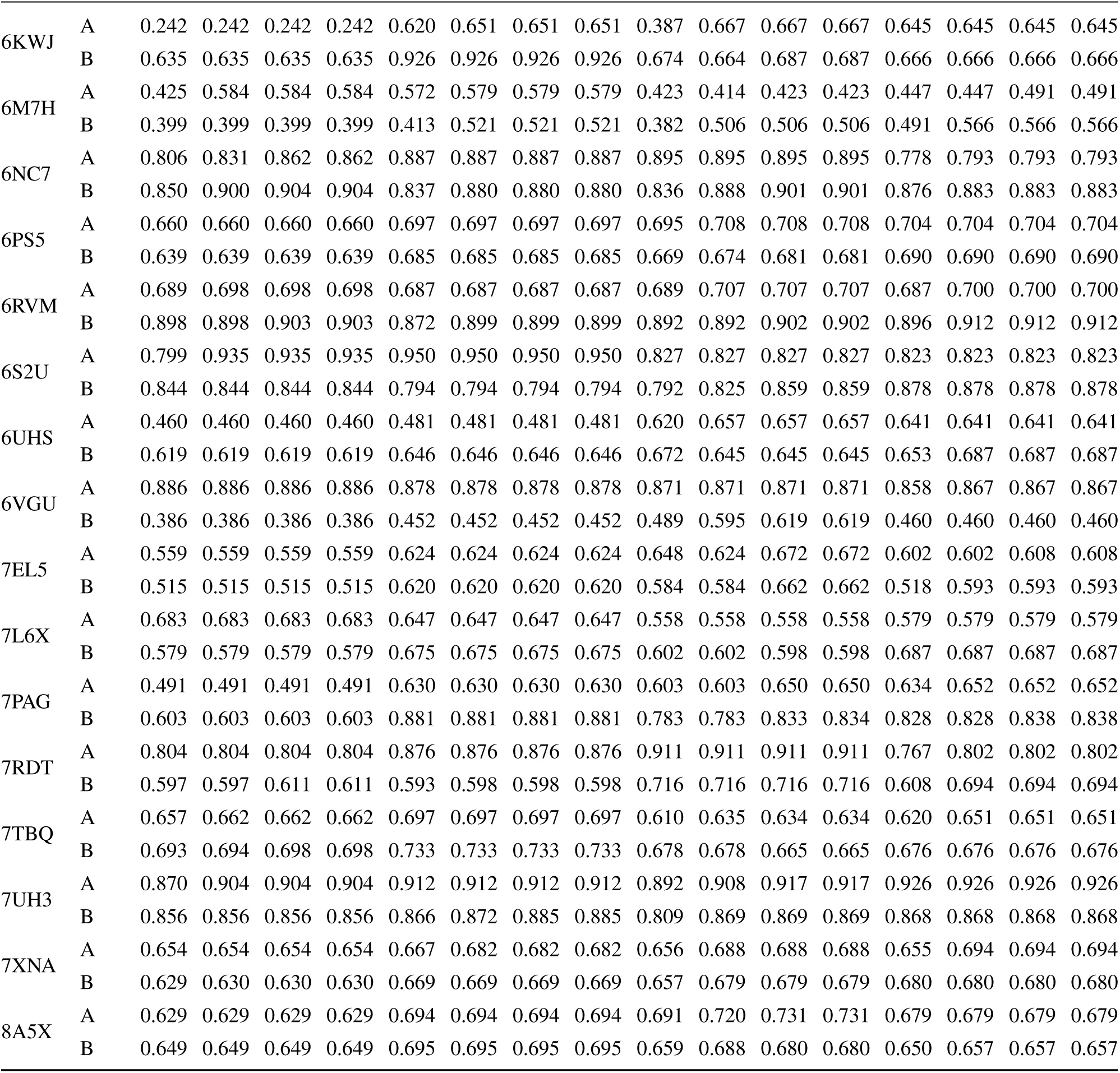
The highest TM-score of representative structures in the fixed-size ensemble to the reference states. For each benchmark protein (row pair), we report the highest TM-scores achieved for reference states A (conf_a) and B (conf_b) by the ensemble of varying sizes (10/20/30/40). This table provides the raw data for Figure 3a.

**Supplementary Table 3.** Clustering yield of predicted structures across methods. For each benchmark protein, the table reports the number of non-redundant conformational clusters (numerator) obtained from the structures generated by the method (denominator) using the same clustering protocol.

| PDB | ProCEDiS (NSGS) | AlphaFlow | BioEmu | PDB | ProCEDiS (NSGS) | AlphaFlow | BioEmu |
| --- | --- | --- | --- | --- | --- | --- | --- |
| 1DQV | 43/220 | 77/250 | 937/10000 | 5AA4 | 45/236 | 4/250 | 553/10000 |
| 1EEH | 7/28 | 2/250 | 1232/10000 | 5AYN | 3/19 | 1/250 | 497/10000 |
| 1FNH | 83/198 | 22/250 | 322/10000 | 5C91 | 8/76 | 3/250 | 410/10000 |
| 1L7C | 25/124 | 7/250 | 1022/10000 | 5CRC | 1/1 | 238/250 | 6953/10000 |
| 1NPR | 61/236 | 106/250 | 1250/10000 | 5IC1 | 37/208 | 10/250 | 1218/10000 |
| 1T3W | 6/37 | 25/250 | 1903/10000 | 5IIT | 42/237 | 28/250 | 1212/10000 |
| 1VS0 | 9/96 | 10/250 | 756/10000 | 5JGL | 6/32 | 4/250 | 63/10000 |
| 1XO5 | 14/122 | 9/250 | 1126/10000 | 5KED | 98/238 | 87/250 | 2358/10000 |
| 2AOB | 44/188 | 9/250 | 308/10000 | 5LB8 | 19/114 | 5/250 | 478/10000 |
| 2B3O | 108/297 | 12/250 | 2161/10000 | 5LE6 | 26/124 | 25/250 | 1546/10000 |
| 2BVN | 4/44 | 2/250 | 165/10000 | 5LEC | 140/308 | 154/250 | 6262/10000 |
| 2C0E | 14/141 | 21/250 | 932/10000 | 5LY6 | 16/157 | 5/250 | 477/10000 |
| 2H2Z | 3/43 | 18/250 | 918/10000 | 5O2K | 8/27 | 2/250 | 170/10000 |
| 2I4I | 11/94 | 4/250 | 765/10000 | 5TZY | 154/300 | 231/250 | 9988/10000 |
| 2QDF | 29/89 | 25/250 | 570/10000 | 5U09 | 1/1 | 167/250 | 820/10000 |
| 2SRC | 79/297 | 10/250 | 845/10000 | 5WBW | 89/288 | 46/250 | 510/10000 |
| 2X28 | 9/121 | 14/250 | 802/10000 | 5XR8 | 45/154 | 191/250 | 9019/10000 |
| 3AJC | 88/306 | 150/250 | 5223/10000 | 6CRF | 231/298 | 8/250 | 4363/10000 |
| 3D8R | 14/53 | 7/250 | 313/10000 | 6GGP | 17/64 | 2/250 | 127/10000 |
| 3FL7 | 42/296 | 54/250 | 3443/10000 | 6HAB | 9/85 | 1/250 | 524/10000 |
| 3KHZ | 7/39 | 12/250 | 3073/10000 | 6IRB | 101/302 | 105/250 | 6906/10000 |
| 3P4Y | 18/108 | 1/250 | 813/10000 | 6IXY | 42/211 | 6/250 | 3282/10000 |
| 3QD9 | 32/228 | 3/250 | 1762/10000 | 6KWJ | 9/75 | 71/250 | 5031/10000 |
| 3TG8 | 9/87 | 10/250 | 126/10000 | 6M7H | 22/43 | 236/250 | 10000/10000 |
| 3USW | 78/284 | 80/250 | 2088/10000 | 6NC7 | 4/33 | 1/250 | 37/10000 |
| 3VKI | 19/134 | 34/250 | 783/10000 | 6PS5 | 57/261 | 225/250 | 9968/10000 |
| 3ZSF | 6/44 | 2/250 | 758/10000 | 6RVM | 2/66 | 1/250 | 144/10000 |
| 4A3P | 21/151 | 10/250 | 698/10000 | 6S2U | 9/53 | 3/250 | 240/10000 |
| 4AVB | 7/90 | 21/250 | 618/10000 | 6UHS | 18/71 | 129/250 | 7666/10000 |
| 4E50 | 49/204 | 185/250 | 8910/10000 | 6VGU | 120/300 | 8/250 | 584/10000 |
| 4EL1 | 59/273 | 111/250 | 1609/10000 | 7EL5 | 50/253 | 91/250 | 2815/10000 |
| 4JZ3 | 13/20 | 4/250 | 672/10000 | 7L6X | 72/298 | 23/250 | 914/10000 |
| 4LAV | 49/208 | 5/250 | 327/10000 | 7PAG | 58/266 | 2/250 | 163/10000 |
| 4N59 | 1/1 | 250/250 | 10000/10000 | 7RDT | 25/78 | 83/250 | 1174/10000 |

| **PDB** | **ProCEDiS (NSGS)** | **AlphaFlow** | **BioEmu** | **PDB** | **ProCEDiS (NSGS)** | **AlphaFlow** | **BioEmu** |
| --- | --- | --- | --- | --- | --- | --- | --- |
| 4ONA | 34/259 | 85/250 | 1647/10000 | 7TBQ | 65/303 | 250/250 | 10000/10000 |
| 4P0I | 7/34 | 4/250 | 628/10000 | 7UH3 | 5/21 | 5/250 | 31/10000 |
| 4TVY | 36/156 | 1/250 | 872/10000 | 7XNA | 44/246 | 250/250 | 10000/10000 |
| 4X8H | 27/58 | 6/250 | 1116/10000 | 8A5X | 7/22 | 188/250 | 9598/10000 |
| 4XZ0 | 36/273 | 13/250 | 910/10000 |  |  |  |  |

**Supplementary Table 4.**
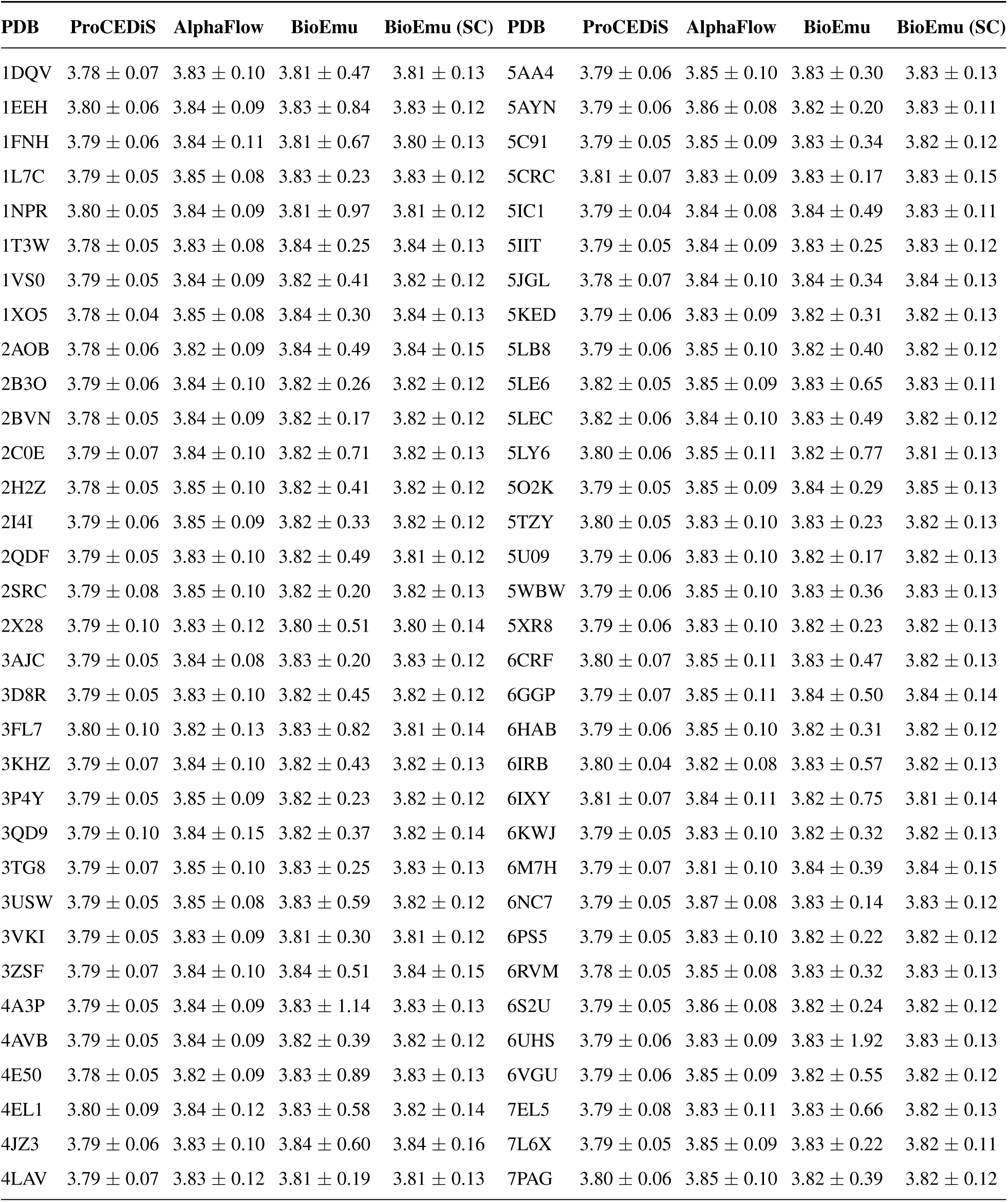
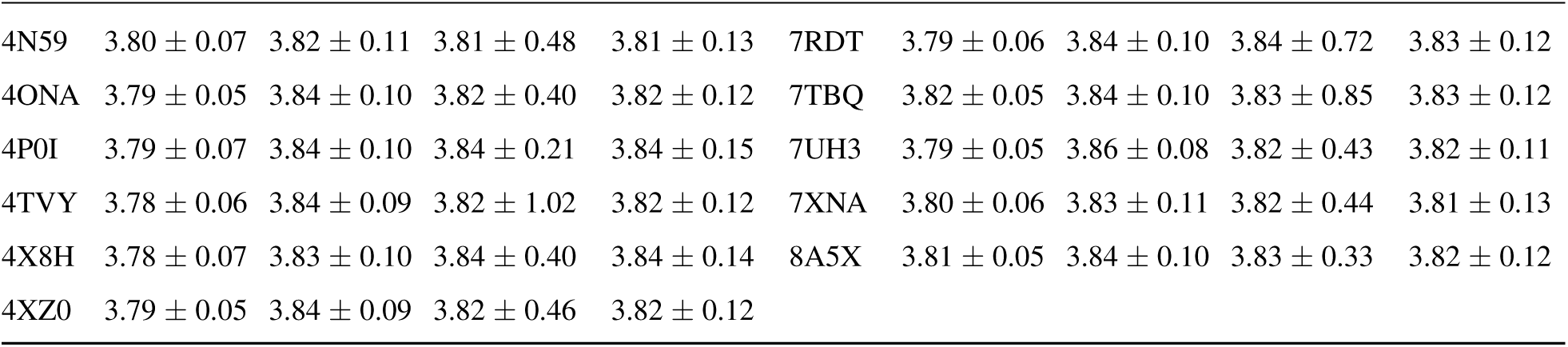
Backbone geometry sanity check viaC*_α_*-C*_α_* distances. Mean standard-deviation (Å) of the C*_α_*-C*_α_* distance between consecutive residues, aggregated over all generated structures per target for each method. This statistic summarizes backbone bond-geometry consistency, where lower degree of variation indicates more regular local geometry (BioEmu (SC) denotes a subset of BioEmu samples after side-chain reconstruction performed following the BioEmu-recommended bioemu.sidechain_relax workflow; no MD equilibration was applied).

**Supplementary Table 5.**
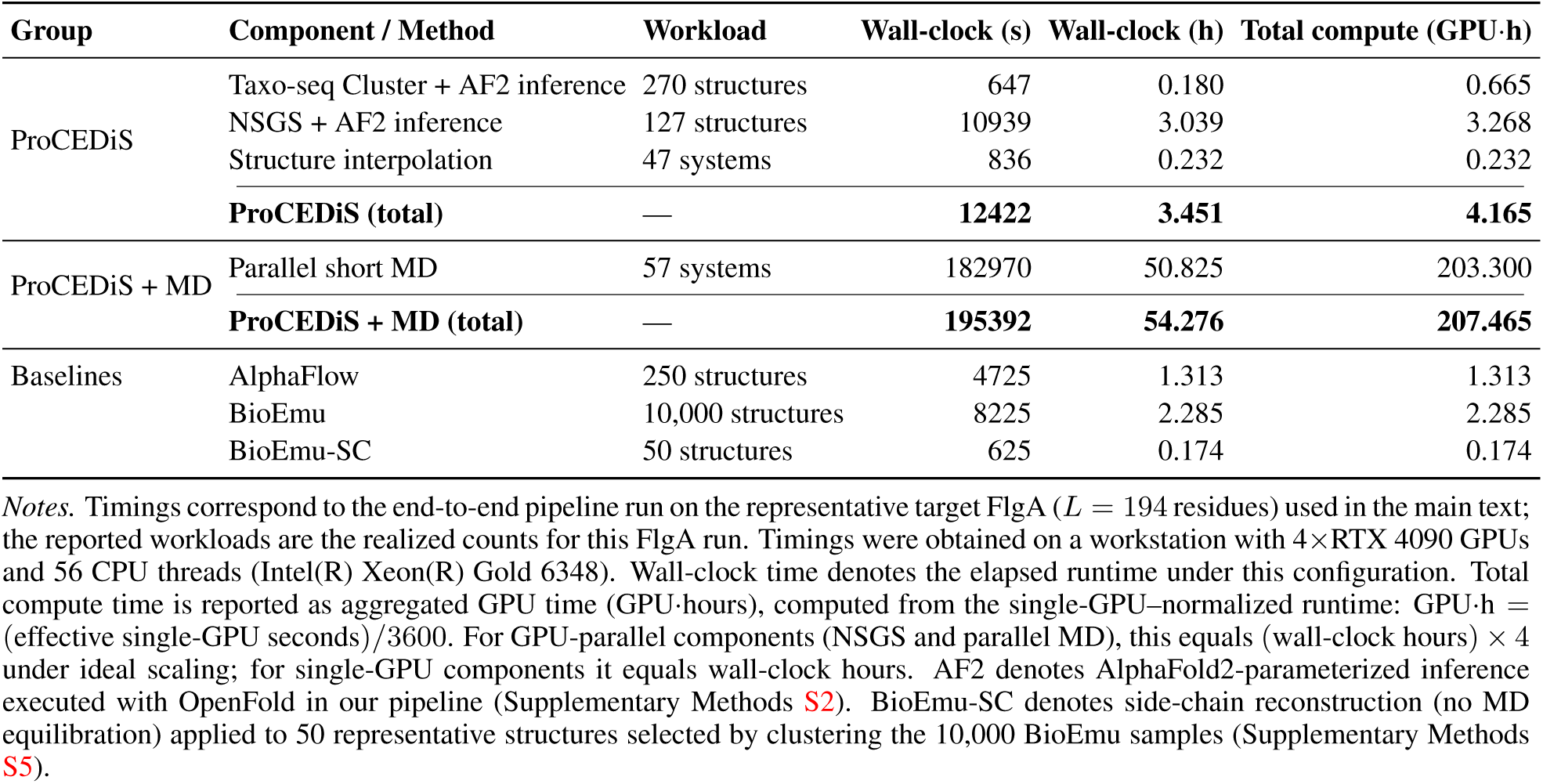
Execution time breakdown across methods. Wall-clock times were measured on a 4×RTX 4090 workstation; total compute time is reported as aggregated GPU time (GPU·hours).

| Group | Component / Method | Workload | Wall-clock (s) | Wall-clock (h) | Total compute (GPU·h) |
| --- | --- | --- | --- | --- | --- |
| ProCEDiS | Taxo-seq Cluster + AF2 inference | 270 structures | 647 | 0.180 | 0.665 |
|  | NSGS + AF2 inference | 127 structures | 10939 | 3.039 | 3.268 |
|  | Structure interpolation | 47 systems | 836 | 0.232 | 0.232 |
|  | **ProCEDiS (total)** | — | **12422** | **3.451** | **4.165** |
| ProCEDiS + MD | Parallel short MD | 57 systems | 182970 | 50.825 | 203.300 |
|  | **ProCEDiS + MD (total)** | — | **195392** | **54.276** | **207.465** |
| Baselines | AlphaFlow | 250 structures | 4725 | 1.313 | 1.313 |
|  | BioEmu | 10,000 structures | 8225 | 2.285 | 2.285 |
|  | BioEmu-SC | 50 structures | 625 | 0.174 | 0.174 |

*Notes.* Timings correspond to the end-to-end pipeline run on the representative target FlgA ( $L = 194$ residues) used in the main text; the reported workloads are the realized counts for this FlgA run. Timings were obtained on a workstation with 4×RTX 4090 GPUs and 56 CPU threads (Intel(R) Xeon(R) Gold 6348). Wall-clock time denotes the elapsed runtime under this configuration. Total compute time is reported as aggregated GPU time (GPU·hours), computed from the single-GPU-normalized runtime: GPU·h = (effective single-GPU seconds)/3600. For GPU-parallel components (NSGS and parallel MD), this equals (wall-clock hours) × 4 under ideal scaling; for single-GPU components it equals wall-clock hours. AF2 denotes AlphaFold2-parameterized inference executed with OpenFold in our pipeline (Supplementary Methods S2). BioEmu-SC denotes side-chain reconstruction (no MD equilibration) applied to 50 representative structures selected by clustering the 10,000 BioEmu samples (Supplementary Methods S5).

### Supplementary Figures

**Supplementary Figure 1.**
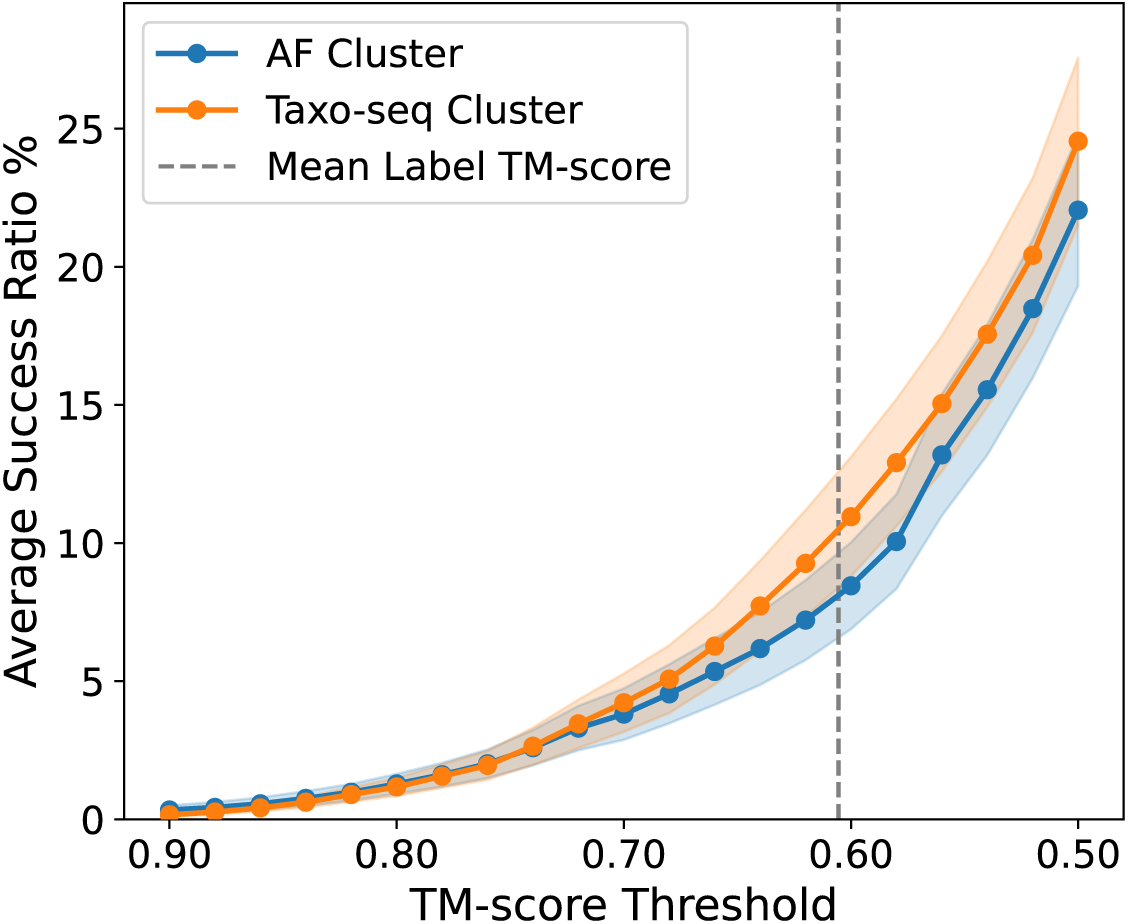
Hit-based success ratio as a function of the TM-score threshold. For each target and TM-score threshold *t*, we compute the fraction of all (predicted structure, reference state) pairs with TM-score ≥ *t* (*i.e.* the average hit number over the two reference states divided by the number of predicted structures; see Supplementary Methods S11). This plot compares the results of Taxo-seq Cluster (orange) and AF Cluster (blue) averaged over the 77 protein targets in the benchmark, with shaded bands indicating the s.e.m. (defined in Equation E7). The dashed vertical line marks the mean TM-score (0.6055) between the two reference states in the benchmark, highlighting the limit of structural distinguishability in this test.

**Supplementary Figure 2.**
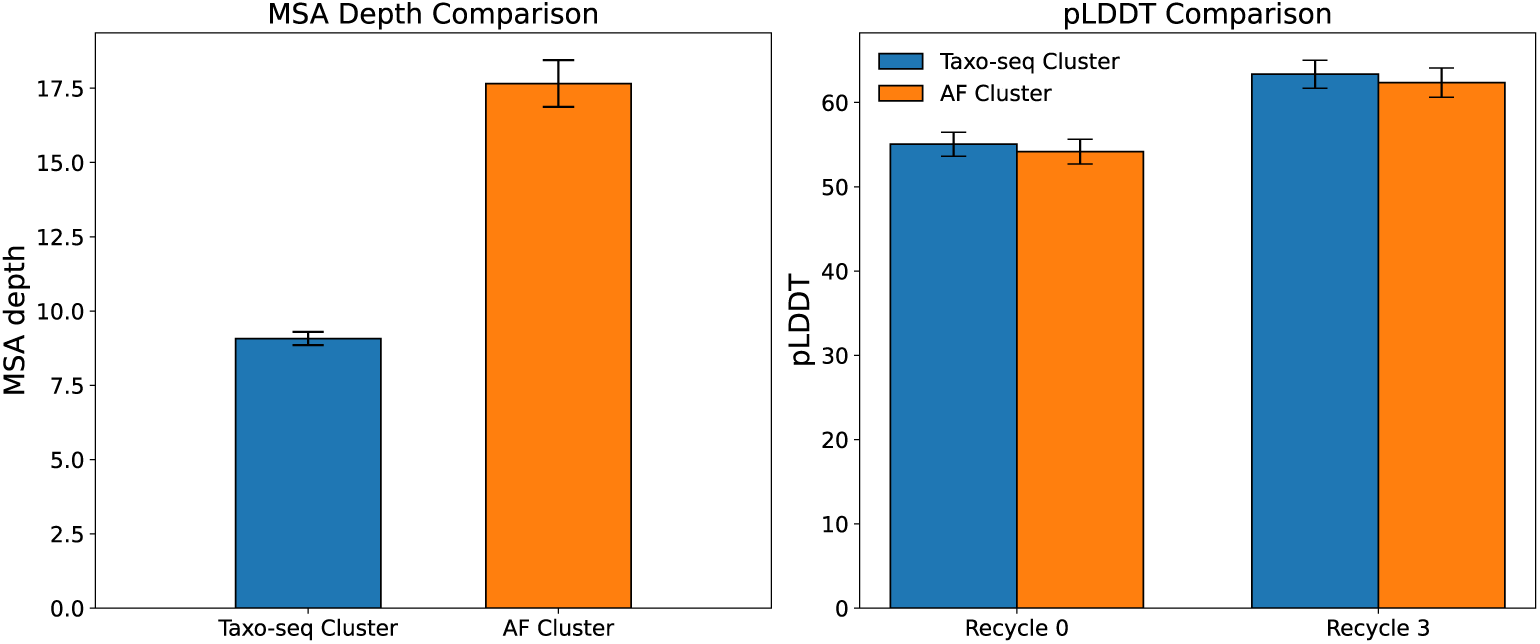
Comparison of Taxo-seq Cluster and AF Cluster in AlphaFold2-based folding. Left panel: the average MSA depth across 77 benchmark targets used for Taxo-seq Cluster (blue) and AF Cluster (orange). Right panel: the mean pLDDT of AlphaFold2-generated structure models across 77 benchmark targets using recycle 0 and recycle 3, respectively, when using Taxo-seq Cluster (blue) and AF Cluster (orange) as the MSA sources. The bar reports the mean across targets, and the error bar indicates the s.e.m. (defined in Equation E7) after excluding NaN values.

**Supplementary Figure 3.**
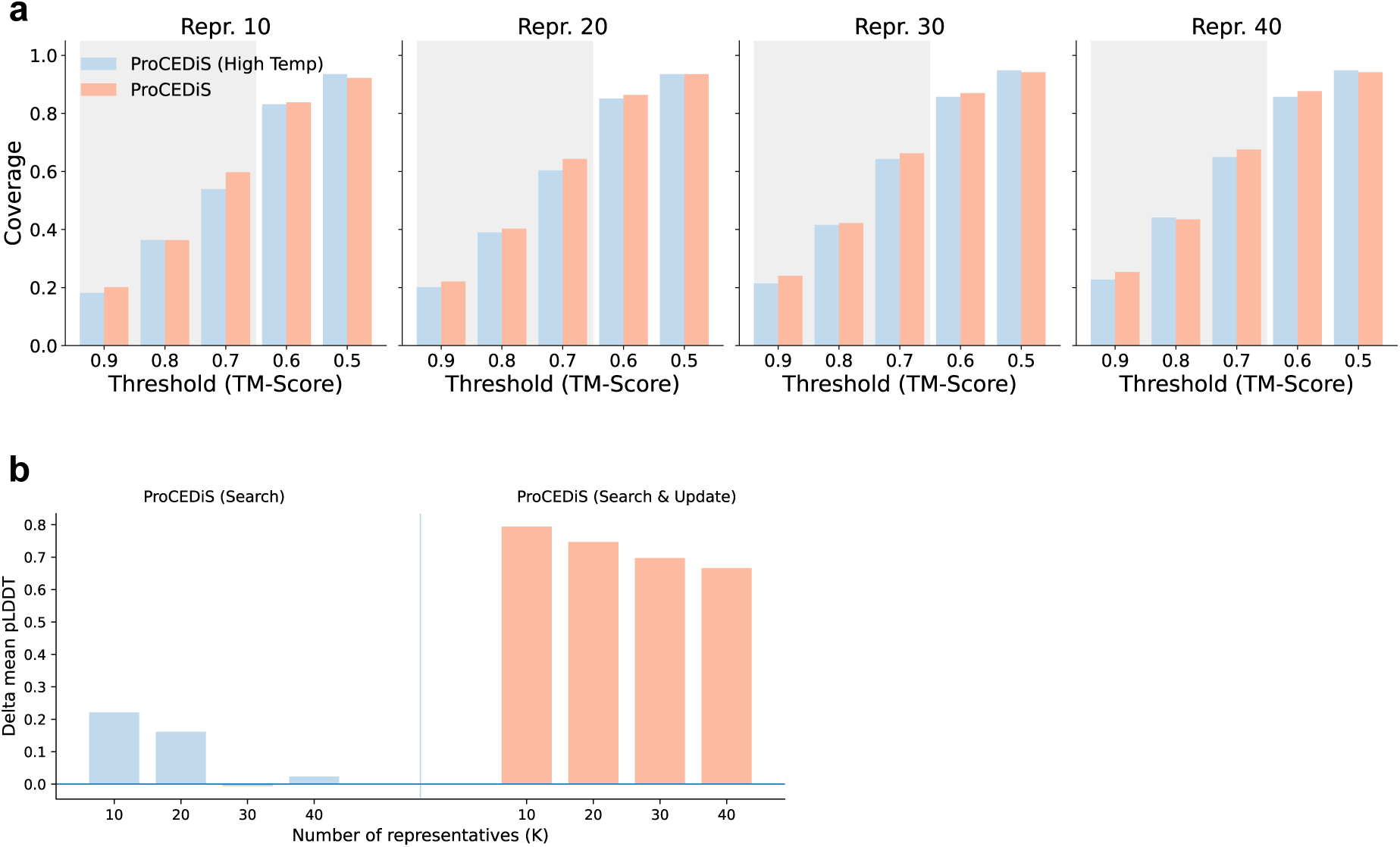
Ablation studies on the neural surrogate. **a)** Evaluation of coverage on the 77 proteins with experimentally determined alternative conformations. This figure is generated following the same protocol as Figure 3a. Here, ProCEDiS (High Temperature) refers to the system in which the action probabilities are nearly uniformly distributed, corresponding to random action choice due to the lack of guidance by the neural surrogate at very high temperatures. In comparison to this control, the formal ProCEDiS pipeline exhibits improved structure coverage, benefited from the effective guidance by our neural surrogate in finding novel structures. **b)** Contribution of the neural surrogate in improving the model quality in the update phase. ProCEDiS (Search) and ProCEDiS (Search & Update) refer to the pipelines without and with the neural-surrogate-assisted structure update phase, respectively. The horizontal axis stands for the number of representative structures in the fixed-size ensemble, while the vertical axis denotes the improvement in modeling confidence (pLDDT) relative to AlphaFold2 predictions based on randomly chosen MSA sub-clusters. Clearly, the neural surrogate makes a positive contribution to the modeling confidence of structures in the final ensemble.

**Supplementary Figure 4.**
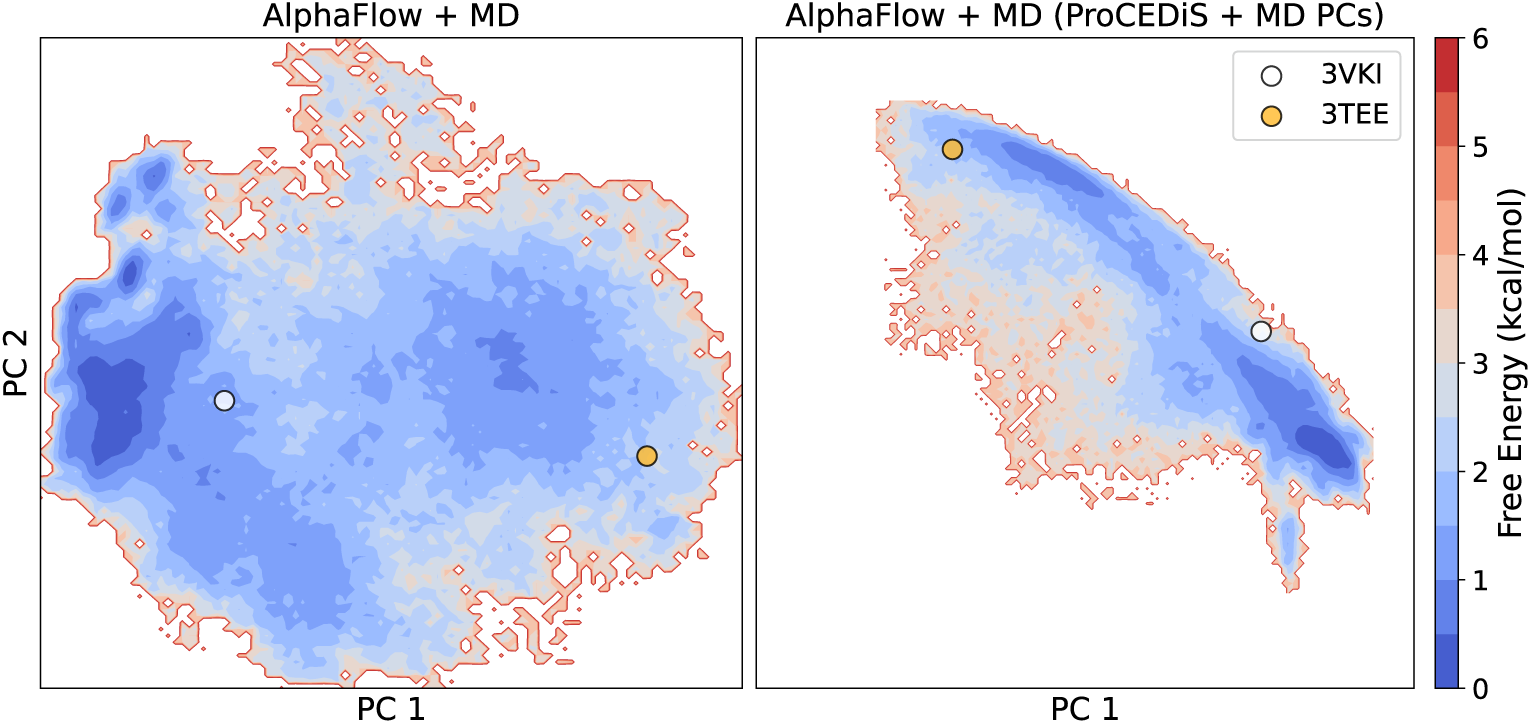
Free energy profile constructed from MD simulations starting from AlphaFlow-generated seeds. The FES is estimated from short-timescale MD simulations initialized from AlphaFlow seeds and is projected into the subspace of the first two PCs. Left: FES in the original projection. Right: the same trajectories re-projected into the PC basis of ProCEDiS + MD, illustrating how the apparent landscape changes with the chosen coordinates. Colored markers indicate projected reference states.

**Supplementary Figure 5.**
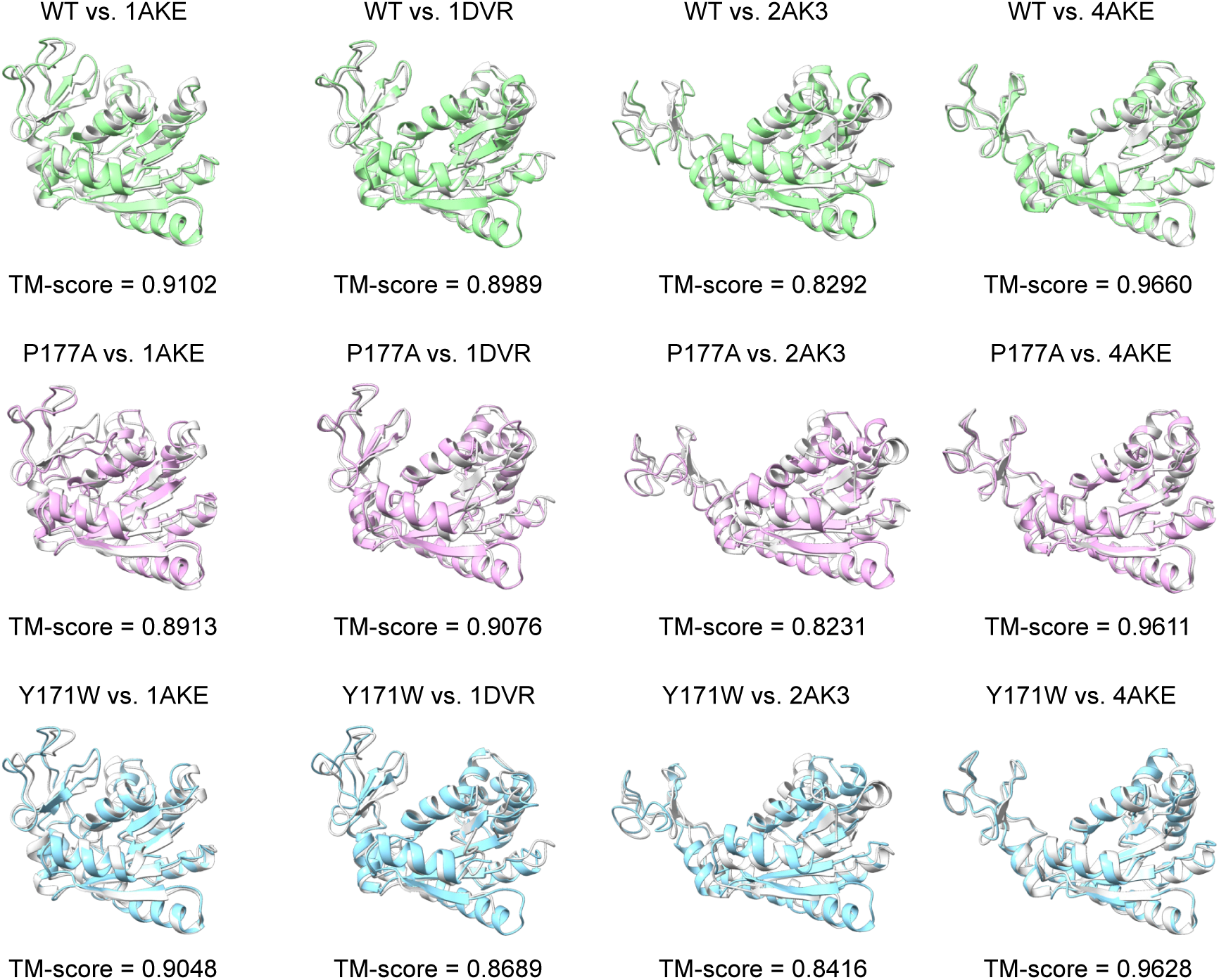
Agreement of structures derived from MD trajectories with known conformational states for ADK variants. Superposition of best sampled structures of ADK against four experimental conformational states (gray) for the wild-type (WT; green), P177A (magenta) and Y171W (blue) ADK, with TM-scores indicated below.

**Supplementary Figure 6.**
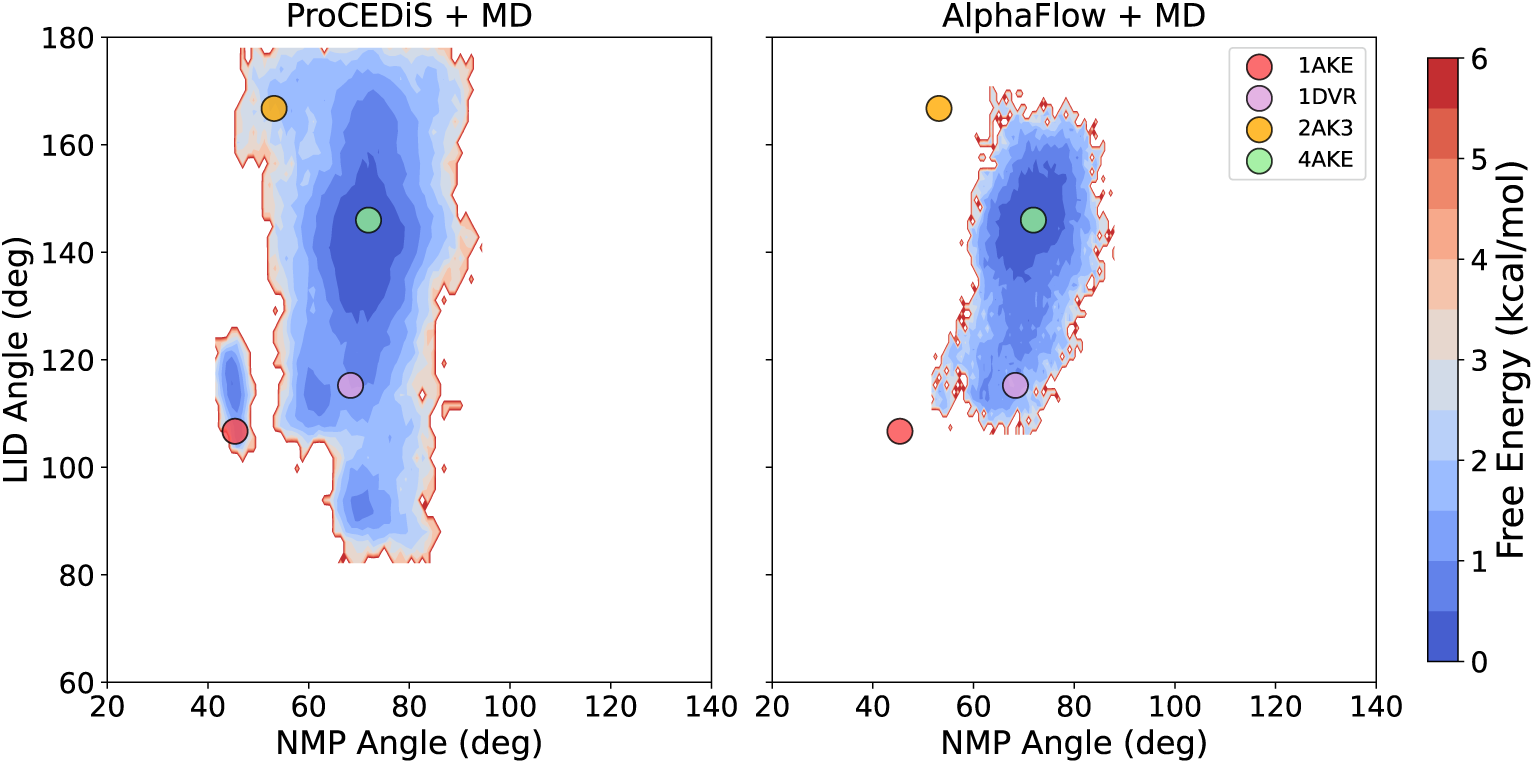
Comparison of free energy landscapes of wild-type ADK between our pipeline and AlphaFlow. MD-based FES of wild-type ADK is projected onto the 2D subspace defined by NMP and LID hinge angles for our pipeline (left) and AlphaFlow (right). Colored markers denote experimentally determined conformations.

**Supplementary Figure 7.**
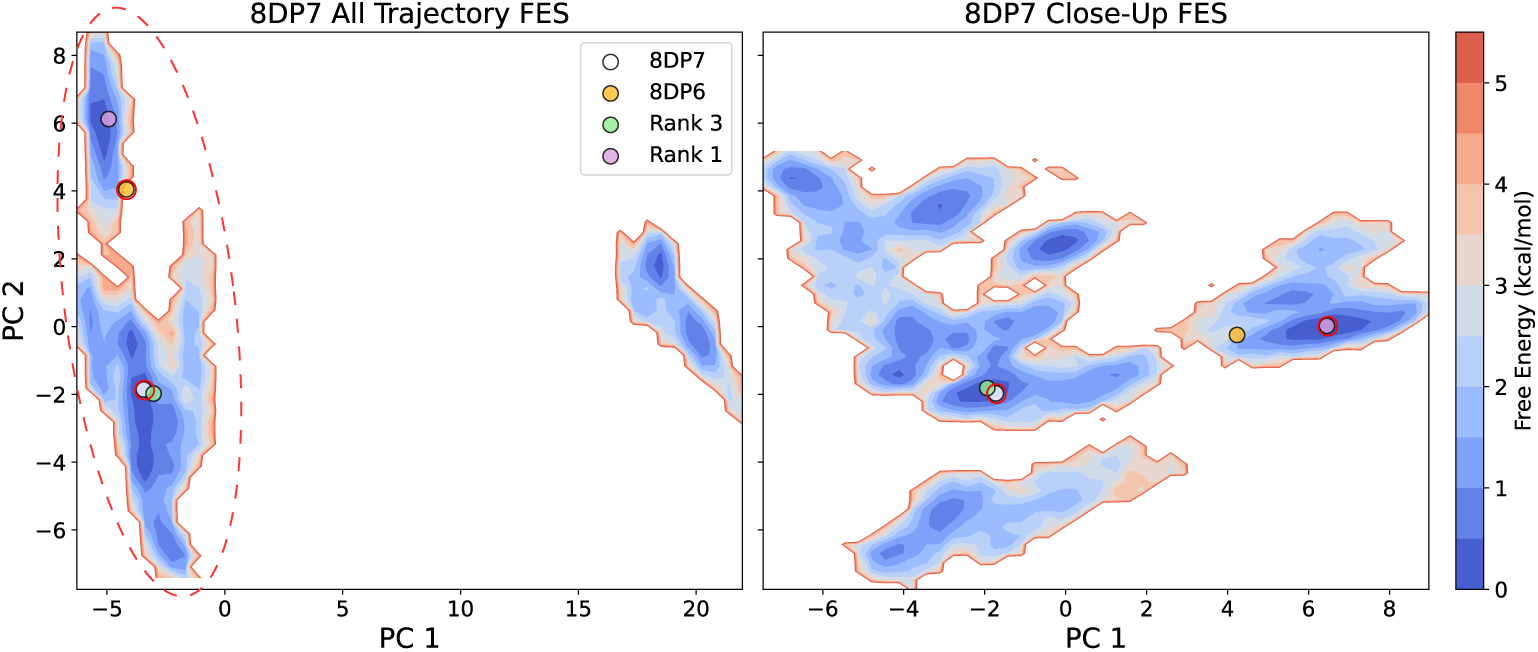
Hierarchical free energy analysis for the 8DP7 system. Left: global PCA projection of conformations sampled by short MD trajectories initiated from ProCEDiS seeds reveals isolated clusters. Right: local PCA performed within a selected region (encircled by the dashed-line ellipsoid) resolves a finer-scale free energy organization and distinctive basins. Experimentally determined reference conformations are indicated by their PDB IDs (8DP7 and 8DP6) and are highlighted by red circles.

**Supplementary Figure 8.**
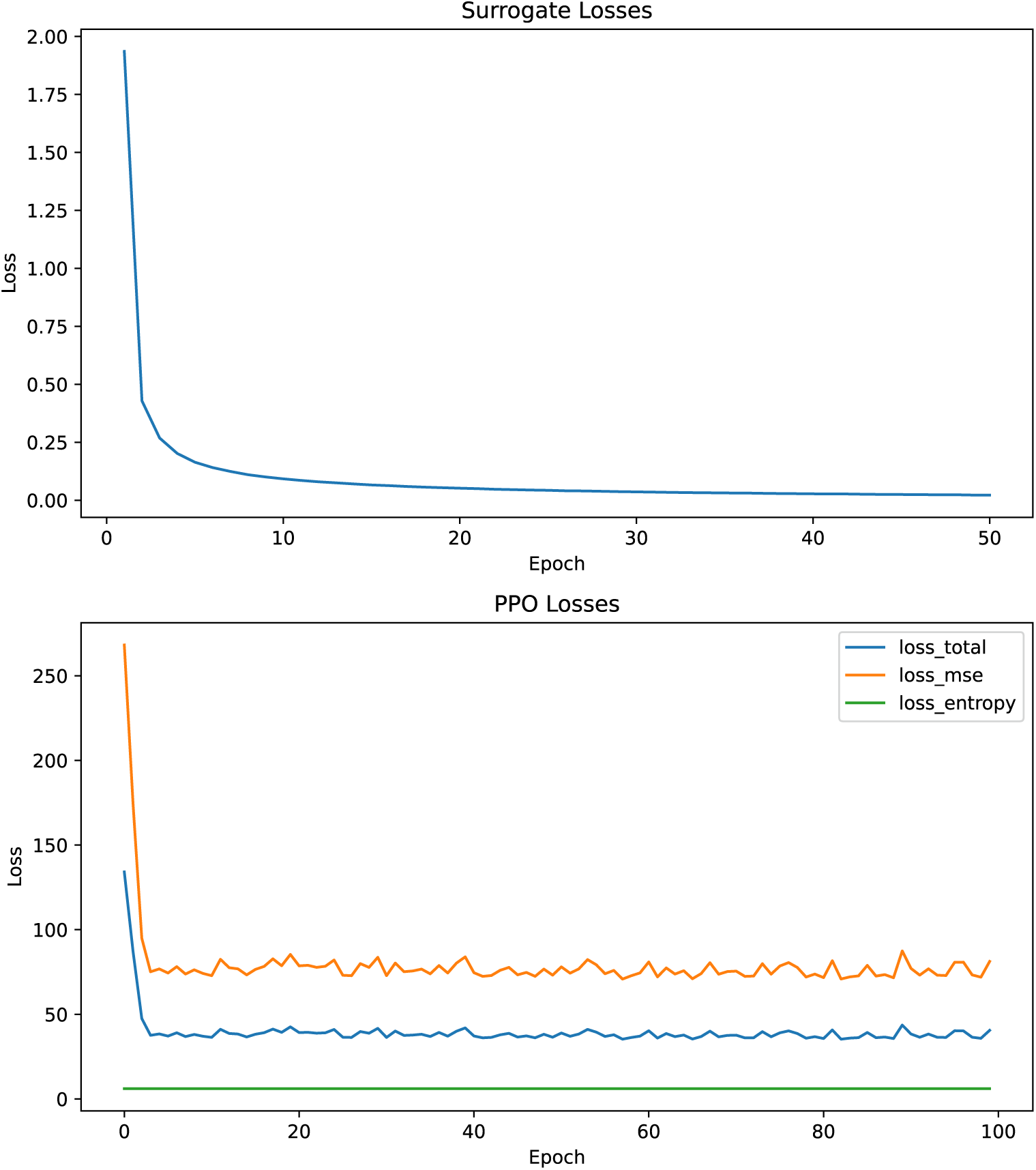
Comparison of training curves in surrogate-guided and RL-based explorations. Top: in our NSGS framework, training loss declines gradually, indicating effective learning by the neural surrogate. Bottom: in the framework implemented with reinforcement learning (RL) using PPO, training loss terms persist after an early drop, negating effective model learning.

